# A neural alternative splicing program controls cellular function and growth in Pancreatic Neuroendocrine tumours

**DOI:** 10.1101/2024.06.13.598849

**Authors:** M. Potiri, C. Moschou, Z Erpapazoglou, G. Rouni, A. Kotsoni, M. Andreadou, M. Dragolia, V. Ntafis, J. Schrader, J. Juan-Mateu, V. Kostourou, S.G. Dedos, M.E. Rogalska, P. Kafasla

## Abstract

Pancreatic neuroendocrine tumours (PanNETs) are a rare heterogeneous group of neoplasms that arise from pancreatic islet cells. The hormone secreting function of pancreatic neuroendocrine cells is altered in PanNETs, rendering these tumours functional or non— functional (secreting excessive or lower levels of hormones, respectively). Genome wide approaches have revealed the genomic landscape of PanNETs but have not shed light on this problematic hormone secretion. In the present work, we show that alternative splicing (AS) deregulation is responsible for changes in the secretory ability of PanNET cells. We reveal a group of alternative microexons that are regulated by the RNA binding protein SRRM3 and are preferentially included in mRNAs in PanNET cells, where SRRM3 is also upregulated. These microexons are part of a larger neural program regulated by SRRM3. We show that their inclusion gives rise to protein isoforms that change stimulus-induced secretory vesicles and their trafficking in PanNET cells. Moreover, the increased inclusion of these microexons results in an enhanced neuronal component in PanNET tumours. Using knock-down and splicing switching oligonucleotides in cellular and animal PanNET models, we show that decrease of the SRRM3 levels or even of the inclusion levels of the three most deregulated microexons can significantly alter the PanNET cell characteristics. Collectively, our study links secretory impairment and nerve dependency to alternative splicing deregulation in PanNETs, providing promising therapeutic targets for PanNET treatment.

## Introduction

Alternative Splicing (AS) of the pre-mRNA generates diverse mRNA and protein isoforms from a single gene. Transcripts from over 95% of human multiexon genes undergo AS, and most of the resulting mRNA splice variants are variably expressed between different cell and tissue types^1,2^. AS is performed by the spliceosome, a large multimeric ribonucleoprotein (RNP) complex, with core-splicing factors necessary for its function and non-core factors regulating the AS decisions. The assembly of spliceosomes at 5′ and 3′ splice sites of the pre-mRNA is typically regulated by RNA-binding proteins (RBPs) that recognize proximal cis-elements, referred to as exonic/intronic splicing enhancers and silencers^3^. Many cell– or tissue-specific and developmentally-regulated AS events are co-ordinately controlled by individual RBPs, and these events are significantly enriched in genes that operate in common biological processes and pathways^4^.

AS is critical for diverse cellular processes, including cell differentiation and development as well as cell reprogramming and tissue remodelling. In almost all cancers, the expression of spliceosome components is altered, contributing to the generation of AS isoforms that are advantageous for tumour development^5–7^. Notably, the impact of the generation of these new isoforms is apparent in almost every hallmark of tumour progression, including cell invasion, angiogenesis, cell metabolism. These changes also significantly affect responses to anticancer therapy and the development of drug resistance. Both the dysregulation of developmental splice site switches and the generation of cancer-specific splicing isoforms contribute to transcriptome changes specific for tumour biology, and these changes are linked to alterations in the activity of splicing factors and regulators.

Many studies have described cancer-associated mutations that disrupt existing splice sites, introduce novel ones or disrupt splicing regulatory elements in exons and introns^5,6^. Antisense oligonucleotides (ASOs) that prevent recognition of these novel elements by spliceosomal regulatory factors have emerged as an entirely novel therapeutic approach to splicing modulation that can alter the AS outcome for a specific pre-mRNA. Preclinical work suggests that ASOs might also have therapeutic value in oncology. However, to reach the point of using these or other tools, one must identify in detail and understand the functions of splice variants that lead to tumour progression and development.

Pancreatic neuroendocrine tumours (PanNETs) are a rare heterogeneous group of epithelial neoplasms with neuroendocrine differentiation that arise from pancreatic islet cells^8^. PanNETs vary in their histological features and clinical symptoms and may be classified as either functional or non-functional, depending on their ability to secrete biologically active hormones and thus elicit characteristic symptomatology. Clinical challenges in their management include – but are not limited to – the inability to establish an early diagnosis in non-functional PanNETs and to accurately localize the source of hormonal secretion in functional ones^8^. Most cases are sporadic, whereas 10% are associated with genetic syndromes, such as Multiple Endocrine Neoplasia type 1 (*MEN1* mutations). The management of PNETs is quite challenging due to the lack of specific biomarkers and efficient treatments, and long-term progression-free survival for patients is rare^9,10^.

During the last years, there has been information on selected splicing factors that show altered expression levels in PanNETs and thus affect tumour aggressiveness or growth potential^11,12^. However, there is a lack of systematic analyses and identification of disease related splicing variants that affect PanNET progression and thus could be explored for therapy development. In the present study, we used an established RNA-seq pipeline to thoroughly characterise the AS changes that take place in PanNET progression. The levels of AS-derived mRNA isoforms were compared between normal human islets and PanNET patient samples. To nominate splicing networks mis-regulated in PanNET tumours, we correlated the expression levels of genes coding for splicing factors with the altered AS events. We identified a group of neural cell specific splicing factors that are potential mediators of the AS deregulation. Our bioinformatic analyses singled out SRRM3 as a splicing regulator critical for the PanNET characteristic AS changes. Use of *in vitro* and *in vivo* models allowed us to verify the detected AS deregulation in PanNETs and show their significance for neuroendocrine cell function and tumour growth. Thus, we uncover the mechanistic details of a new neural specific splicing program that increases the neural component of neuroendocrine cells and drives tumour growth.

## Methods

### Alternative splicing and Differential gene expression analyses

Access was gained to publicly available RNA sequencing datasets from normal pancreas and from PanNET patients. In detail, two datasets were analyzed as normal pancreatic tissue and two as PanNET. PanNET public RNA sequencing datasets can be found under accession numbers EGAS00001001732 and GSE118014^13,14^. Normal pancreatic tissue datasets derived from VastDB repository for each different type of pancreatic cell and under accession number GSE50398^15^ for whole pancreatic islets. At the same time, a publicly-available human adenocarcinoma dataset was also analyzed for splicing to use for comparisons with PanNET samples, with accession number GSE79668^16^. FASTQ reads were aligned with the human VASTDB library (hg38, vastdb.hs2.23.06.20).

Alternative splicing analysis was performed with VAST-TOOLS v2.5.1^15^ and results were depicted as changes in percent-spliced-in values (ΔPSI). PSI values for single sample replicates were quantified for all types of alternative events, comprising single and complex exon skipping events (S, C1, C2, C3, ANN), microexons (MIC), alternative 5’ and 3’ splice sites (Alt5 and Alt3, respectively) and retained introns (IR-S, IR-C). Events showing splicing change |ΔPSI|> 15 and P-Adjusted value < 0.05 calculated using non-parametric Kruskal-Wallis test (R library matrixTests, v0.2), followed by Dunn’s Test (R library rstatix, v0.7.2), were considered preferred regulated events under tumoral conditions.

For differential gene expression analysis, FASTQ reads were aligned to the GRCh38 human reference genome assembly using the STAR alignment tool (vSTAR-2.7.10a)^17^. The aligned reads were then used to generate BAM files and extract the gene reads for downstream analyses. iDEP workflow was used to generate Log_2_FoldChange values between control and PanNET samples^18^. Genes showing |Log_2_FoldChange| >0.58 and P-Adjusted value <0.05 were considered differentially expressed.

### ORF impact prediction

Potential ORF impact of alternative exons was predicted as described by Irimia et al^19^. Exons were mapped on the coding sequence (CDS) or 5’/3’ untranslated regions (UTR) of genes. Events mapping on the CDS were divided into CDS-preserving or CDS-disrupting.

### Cell markers

Utilizing the PanglaoDB database^20^, specific cell markers were selected and downloaded regarding normal pancreatic tissue, including alpha, beta, pancreatic progenitor, stellate and peri-islet Schwann cells. The retrieved cell marker data was used to generate a heat-map plot illustrating the expression patterns across our samples according to the color scale employed. Visualization was made using pheatmap (v1.0.12) package in R, with rows representing cell types and columns representing our samples.

### RNA maps analysis

To generate the RNA maps, we used the rna_maps function (Matt software v1.3.0^21^), using sliding windows of 11 nucleotides. Searches were restricted to the affected exons and the first 75 nucleotides of the upstream intron. Regular expression was used to search for the binding motif of SRRM3 (UGC). RNA maps for the SRRM3 motif were analyzed. Cassette exons were grouped as follows: up ΔPSI >15 and PSI margin between groups >5, down ΔPSI < –15 and PSI margin between groups >5. The sequence of first 15 nts of exons and the first 75nt of introns (sliding window = 11, p value ≤ 0.05 with 1000 permutations) were compared with the non-changing exons (ndiff –2>ΔPSI >2 and average PSI controls < 95 and ΔPSI ≤ 5).

### Gene Ontology analysis

Ensembl gene IDs for the significant MIC splicing events (Length <50) were analyzed using clusterProfiler (v4.2.1) R package. Statistical significance was defined with p-Adjusted value< 0.05 in the comparison of PanNET to islets and the positive false discovery rate with q value<0.05. The obtained GO annotations for our genes were further reduced using revigo R library (rrvgo, v1.8.0), to identify enriched functional categories.

### Correlation analyses

For comparison with cancer samples from the Cancer Genome Atlas Program (TCGA), lists of significantly deregulated splicing events between tumors and normal tissues (|ΔPSI|>5, P value < 0.05) were obtained from the analysis done by Head et al (2021)^22^. Using Pearson coefficient method and the corrplot R library (v0.92), the association between altered exonic splicing events in PanNET in comparison to islets (|ΔPSI|>15, P-Adjusted value < 0.05) and deregulated events in other types of cancer, was evaluated. For the correlation thresholds, |association value|>0.6 and FDR<0.05 were applied.

For the correlation between splicing factors and splicing events, again the corrplot R library with Pearson coefficient was used. The splicing factors with |Log2FoldChange|>0.58 and P-Adjusted value <0.05 emerging from the differential expression analysis, were associated with the deregulated exonic splicing events in PanNET compared to islets (|ΔPSI|>15, P-Adjusted value <0.05), applying fdr<0.05.

### SRRM3 gene expression plots

SRRM3 log_2_FoldChange values were acquired from Head et al. (2021)^22^ and visualized in R, using ggpubr library (v0.6.0) in comparison to PanNET. In addition, individual cRPKM values for SRRM3 gene were downloaded from VastDB repository for distinct types of normal tissues, including cerebellum, retina, whole brain, cortex, adrenal, heart, endometrium, prostate, adipose, bladder, thyroid, breast, melanocytes, lung, kidney, endothelium, muscle and liver, to compare them with SRRM3 expression in PanNET and the normal pancreatic tissues we analyzed.

### Statistical analyses and plots

Statistical tests were performed as indicated in figures legends using R (v4.2.1)^23^ or GraphPad Prism (version 8). Principal Component Analysis (PCA) was performed on PSI values of the top 5000 most variable events and visualized using ggfortify (v0.4.16) and ggplot2 (v3.4.2) packages in R. Venn diagram was plotted using VennDiagram package in R (v1.7.3). For heatmaps PSI were plotted using pheatmap (v1.0.12) package in R. Hierarchical clustering was performed using a Ward’s method and Euclidean distance as the distance metric.

### Cell Culture

Human pancreatic cancer cell lines QGP-1 and NT-3 were used for this study. QGP-1 (RRID:CVCL_3143) is a neuroendocrine tumor cell line of pancreatic somatostatinoma origin. QGP-1 were cultured in RPMI1640 medium (GIBCO Cat# 31870-025) supplemented with 10% fetal bovine serum (FBS), 1% L-Glutamine+ and 1% penicillin–streptomycin. The NT-3 (RRID:CVCL_VG81) cell line originated from a pancreatic neuroendocrine tumor G2 stage, were cultured under semi-adherent conditions in collagen IV-coated plates (Sigma C7521-5MG, final concentration 50μg/ml) in RPMI 1640 GlutaMAX™ Supplement (Gibco/Thermofisher, Cat#61870010) accompanied with 10% FBS, penicillin/streptomycin, epidermal growth factor (EGF, 20 ng/mL), and fibroblast growth factor 2 (FGF2, 10 ng/mL). The cells were maintained in a standard humidified incubator at 37°C in a 5% CO_2_ atmosphere.

### RIP1-Tag2 mouse model

The mouse model RIP1-Tag2 was used for validation of the selected alternative splicing events. Three timepoints (T1, T2, T3) of mice growth were used. RIP1-Tag2 mice start to develop beta-cell hyperplasia at around 8 weeks of growth (T1) and they normally die between 12 and 14 weeks of age (T2). Upon high-carbohydrate diet, they can survive until 17-18 weeks along with developing highly invasive carcinomas (T3)^24^. The mice were euthanized and either islets were extracted for RNA or tumors/total pancreases were dissected for RNA isolation and paraffin embedding for subsequent analyses.

In total, 25 wild type (WT) and 23 transgenic (RT2) mice were used for validations. The study was approved by the local project evaluation committee (License number 278206-01/04/2022). Animals were maintained in a climate-controlled environment at 21-22°C, 50-60% humidity, with a light-dark cycle (12:12 h), standard laboratory chow and 5% sugar water abundance ad libitum.

### RNA isolation, cDNA synthesis and PCR/qPCR

Total RNA from murine pancreas was extracted using RNeasy Mini Kit (Qiagen) following the manufacturer’s instructions. RNA isolation from cell lines was performed with the TRIzol® reagent (Thermo Fisher Scientific, Inc). Islet isolation and subsequent RNA extraction was carried out as described by Li et al. ^25^.

Reverse transcription (RT) was carried out with 400 ng to 1μg of total RNA, using random hexamer primers (Invitrogen), OligodT (New England BioLabs, Inc.), RNaseOUT^TM^ Recombinant Ribonuclease Inhibitor (Thermo Fisher Scientific) and Protoscript II reverse transcriptase (New England BioLabs, Inc.), according to manufacturer’s instructions.

Semi-quantitative PCR reactions were carried out with 2 µl of cDNA, with different cycling and temperature parameters for each selected alternative splicing event (Table S5). Inclusion/exclusion bands for these events were analyzed by acrylamide electrophoresis and quantified using ImageJ (v1.53t). Quantitative PCR (qPCR) was performed in a Corbett Rotor-Gene™ RG 6000 using Kapa SYBR®FAST (KAPA Biosystems) qPCR mix.

### Mouse Xenografts study

For this study, 1×10^6^ cells suspended in 100µl of PBS:Matrigel (1:1; Corning) of QGP-1 NT and QGP-1 Sh1 or Sh2 cells, were injected into the left and right flank of 15 week-old NOD-SKID mice (NOD.CB17-Prkdcscid/J, Charles River, Strain code: 634) respectively. A total of 9 mice were used and 3D ultrasound images (via 3D scans) were acquired using the Vevo 3100 Imaging System (FUJIFILM VisualSonics). Tumor growth was monitored for up to 4 weeks and weekly calculations were carried out using the VevoLAB software (version 5.8.1.3266). At the endpoint, mice were euthanized, and both tumors and pancreases from each mouse were weighed and collected for RNA isolation and paraffin embedding for subsequent analyses. The study was approved by the local project evaluation committee (License number 278206-01/04/2022).

### Immunohistochemistry and H&E staining

After pancreases were obtained from RIP1-Tag2 mice or at the end-point of the xenograft experiment, parts of the pancreases or tumors were dissected, fixed in formalin and embedded in paraffin. Sections were cut at 5 μm thickness, deparaffinized, re-hydrated in descending ethanol grades, washed and boiled at 95°C for 15 minutes in sodium citrate buffer pH 6.0 for antigen retrieval. Blocking was performed with 5% BSA for 1 hr and sections were then incubated with the following primary antibodies overnight at 4°C diluted in BSA: Insulin (Cat #EPR17359, 1:200), TBB3 (Cat #EP1569Y,1:200). Sections were subsequently washed and incubated in 3% H_2_O_2_ solution for 10 minutes. HRP-conjugated goat anti-rabbit IgG (Cat # sc-2004, 1:5000) was applied for 1h at RT and the DAB Substrate Kit was used to visualize the signal. The sections were counterstained with hematoxylin and images were obtained with a NIKON Eclipse E600 microscope, equipped with a Qcapture camera.

### Transfection of QGP1 and NT-3 (siRNA, shRNA, RINS1 plasmid)

For RNA-mediated interference, cells were transfected with control (scramble) or SRRM3-siRNA (L-016790-02-0005 SMARTPool siRNA) at 30 nM final concentration for 72 h, using the Lipofectamine RNAiMAX transfection reagent (Thermo Fisher Scientific), according to manufacturer’s instructions. Alternatively, NT-3 cells were transfected with control or *SRRM3*-siRNA by electroporation using the Super Electroporator NEPA21 (Nepa Gene Co., Ltd; Poring pulse: pulse voltage 175 V; pulse interval 50 ms; pulse width 5 ms; pulse number 2; and Transfer pulse: pulse voltage 20V; pulse interval 50 ms; pulse width 50 ms; pulse number 5). For co-transfection of NT-3 cells with siRNAs and RINS1 plasmid (gift from Dmytro Yushchenko, Addgene plasmid # 107290, doi: 10.1016/j.chembiol.2017.03.001), lipofectamine 2000 transfection was used (Thermo Fisher Scientific), according to manufacturer’s instructions.

For antisense oligonucleotides (ASOs) transfection, QGP-1 cells were transfected with control (scramble) siRNA or ASO targeting three individual microexons (CADPS2, DADNA1D, PTK2) at 40nM final concentration for 72h using the Lipofectamine RNAiMAX transfection reagent (Thermo Fisher Scientific), according to manufacturer’s instructions. ASOs were designed as described in by Juan-Mateu et al^21^.

For viral transduction in QGP-1 cells, the sequences of shRNA used are: Sh1 (sh targeting SRMM3 CGTTACGAACACAGGAATCTC) and Sh2 (sh targeting SRMM3 GCATAGGATAACATGTGCTTT). Lentiviral vector production and transduction were performed using standard protocols. Individual clones were selected with puromycin (final concentration 0.5μg/ml) and kept in cell culture.

### Intracellular Calcium Measurements

QGP1 and NT3 cells were seeded in 96-well cell culture plates at a confluency of 30000 cells/well and the following day were assayed for [Ca^2+^]_i_ measurements using a FlexStation 3 96-well multi-mode microplate reader (Molecular Devices). Before the assays, cells were loaded with Fluo-4/AM (F14201, ThermoFisher ScFientific) in 100 μL of HEPES-Buffered Saline (HBS:135 mM NaCl, 5.9 mM KCl, 11.6 mM HEPES, 1.5 mM CaCl_2_, 1.2 mM MgCl_2_, 1 mM glucose (pH 7.3) that contained 0.02% Pluronic-F127 (P3000MP, ThermoFisher Scientific, 20% (w/v) solution in DMSO) and supplemented with 0.1% Bovine serum albumin (A9418, Sigma-Aldrich). Cells were incubated for 1 hr at 37°C, then the medium was removed and replaced with HBS, supplemented with 0.1% Bovine serum albumin, for 45 min at 37°C before the assays. Measurements were taken at 25°C on the FlexStation 3 with reads taken at 1.5 sec intervals at Exc./Em. 490/520 nm wavelengths, in FLEX mode using medium photomultiplier tube (PMT) gain and 5 flashes per read. Cells were assayed in a volume of 180 μl before the addition of 20 μL 0.5 M KCl or 20 μL 0.2 M Glucose in each well at 120 sec after the onset of measurements. Assays were carried out for 12 min and then measurements of Fmax and Fmin were taken by the addition of Triton X-100 (10591461, Fisherscientific) (2% final concentration) and BAPTA (196418, Sigma-Aldrich) (10 mM final concentration), sequentially and at 1 min intervals on each well upon completion of measurements. Results were visualized with the SoFtMaxPro^©^ 7 software (Molecular Devices) and fluorescent intensities were converted to [Ca^2+^]_i_ concentrations using the equation [Ca^2+^]_i_ (nM) _=_ K_d_ x (F-Fmn/Fmax-F) where K_d_ for Fluo-4 is 335 nM. When Nifedipine was used, it was added in each well offline 3 min before the start of fluorescence measurements to give a final concentration of 10 μΜ in each well. At least 4 biological replicates were carried out with 6 wells assayed for each condition in each biological replicate. Data on [Ca^2+^]_i_ was visualised in GraphPad Prism (v8) with error bars representing ±SEM.

### MTT cell viability assay

24h after transfection, QGP-1 cells were plated at a density of 5000 or 20000 cells/well in triplicates on a 96-well plate, for nifedipine treatment or siRNA/ASO transfection. At 48h or 72h post-transfection, cell viability was assessed using MTT reagent (Sigma) dissolved in PBS (5mg/ml). On the day of measurement, fresh medium containing diluted MTT (1:10, 0.5mg/ml final concentration) was added and incubated for 3 h at 37 °C. After removing the medium, formazan crystals were dissolved in 200μl solution of DMSO. MTT reduction was quantified by measuring the absorbance at 570 nm using the Tecan Sunrise microplate reader and the Magellan^TM^ software.

### Analysis of insulin secretion with the ratiometric reporter RINS1

The protocol described in Schifferer *et al.*^26^ was used, with small modifications. Briefly, NT-3 cells were co-transfected with control or *SRRM3*-siRNA and RINS1 plasmid. After 72 h, the medium was replaced with low glucose containing Imaging buffer (115 mM NaCl, 1.2 mM CaCl_2_, 1.2 mM MgCl_2_, 1.2 mM K_2_HPO_4_, 0.2% glucose, 20 mM Hepes pH 7.4) and the cells were allowed to adapt for 2 h at 37°C. Insulin secretion was stimulated by exposing cells to Imaging buffer supplemented with 20 mM glucose and 30 mM KCl. Supernatants were collected after 0– and 10-min and cleared by centrifugation at 600 x *g* prior to measurement of sfGFP and mCherry fluorescence intensities. Cells were lysed with lysis buffer (10 mM Tris-HCl pH 7.4, 150 mM NaCl, 0.5 mM EDTA, 0.5% NP-40) supplemented with protease inhibitors, and diluted 3-fold in Imaging buffer prior to fluorescence measurement. sfGFP and mCherry fluorescence intensities were measured on a FlexStation 3 Multi-Mode Microplate reader system and SoftMax Pro 6 software for analysis with the following settings: for sfGFP, Excitation 485 nm; Cut off 515 nm; Emission 520 nm; for mCherry, Excitation 587 nm; Cut off 610 nm; Emission 620 nm.

### Immunostaining with Golgi markers

Cells were fixed for 5 min at –20°C with ice-cold methanol. Rabbit CoraLite® Plus 647-conjugated GOLGA2/GM130 polyclonal antibody (Proteintech, Cat # CL647-11308, 1:100) and mouse beta-tubulin monoclonal antibody (1:100) were used as primary antibodies. Anti-mouse Alexa Fluor 488 (Molecular Probes Cat# A28175) was used as secondary antibody. Analysis of *cis-* Golgi area relatively to cell surface was performed with ImageJ (v1.53t). Analysis of Golgi area relatively to nuclei was performed with Imaris software.

### Western blot analysis

Goat polyclonal anti-GFP and anti-mCherry (SICGEN Antibodies, Cat # AB0020 and AB0040, respectively, 1:1000) were used as primary antibodies. HRP-conjugated donkey anti-goat IgG (Santa Cruz Biotechnology, Cat # sc-2020) was used as secondary antibody.

## Results

### Deregulation of AS differentiates PanNET from normal tissues and other cancer types

To investigate the global changes in mRNA isoform expression between tumours and normal tissue we profiled the AS patterns of publicly available RNAseq datasets of PanNET patients. The general characteristics regarding age, gender, genotype, disease status and grading of the patients are shown in Figure S1A. Most of the samples were derived from non-functional PanNETs and were mostly of low grading (G1 and G2). In parallel, we profiled the AS patterns of publicly available RNAseq from normal human islets^27^ and from normal human pancreatic single cell populations (alpha, beta, acinar and ductal cells^15^. We assessed the expression profiles of neuroendocrine markers for all the samples and thus verified the identity of the normal ones, but also the deregulation of the pancreatic cell markers in PanNET samples (Figure S1B). For each splicing event, the ‘‘percent spliced in’’ (PSI) value was estimated, reflecting the fraction of the mRNA isoforms that are derived from a single gene and include the alternative exon, intron, or splice site. A total of 721551 events, supported by sufficient junction reads to allow their quantification, were identified. Of these events, 239456 were consistently mapped in at least 70 out of 128 samples. The majority of these events involved alternative exons (∼40%), followed by a large number of intron retention events (34%) and smaller numbers of Alt3’ (15%) and Alt5’ (∼10%) (Figure S1C).

To determine the impact of PanNET on the overall AS profiles we employed PCA analyses using the PSI values of the 5000 most variable events (Figure S1D). Distinct grouping of PanNET, islets, alpha/beta and acinar/ductal cells was observed indicative of the different AS pattern in the respective datasets. This grouping was even more apparent when PCA analysis was performed of the PSI values of the most variable alternative exon events (Figure 1A), the type of events that constitutes the majority of mapped events (Figure S1C). Similar PCA analysis of the PSI values of the most variable intron retention events (Figure S1E) did not give distinct grouping for the different tissue/cell categories. Interestingly, the two distinct datasets of PanNET samples that we used^13,14^ clustered very nicely together (Figure 1A). Thus, we focused our analyses on the alternative exon events.

**Figure S1.**
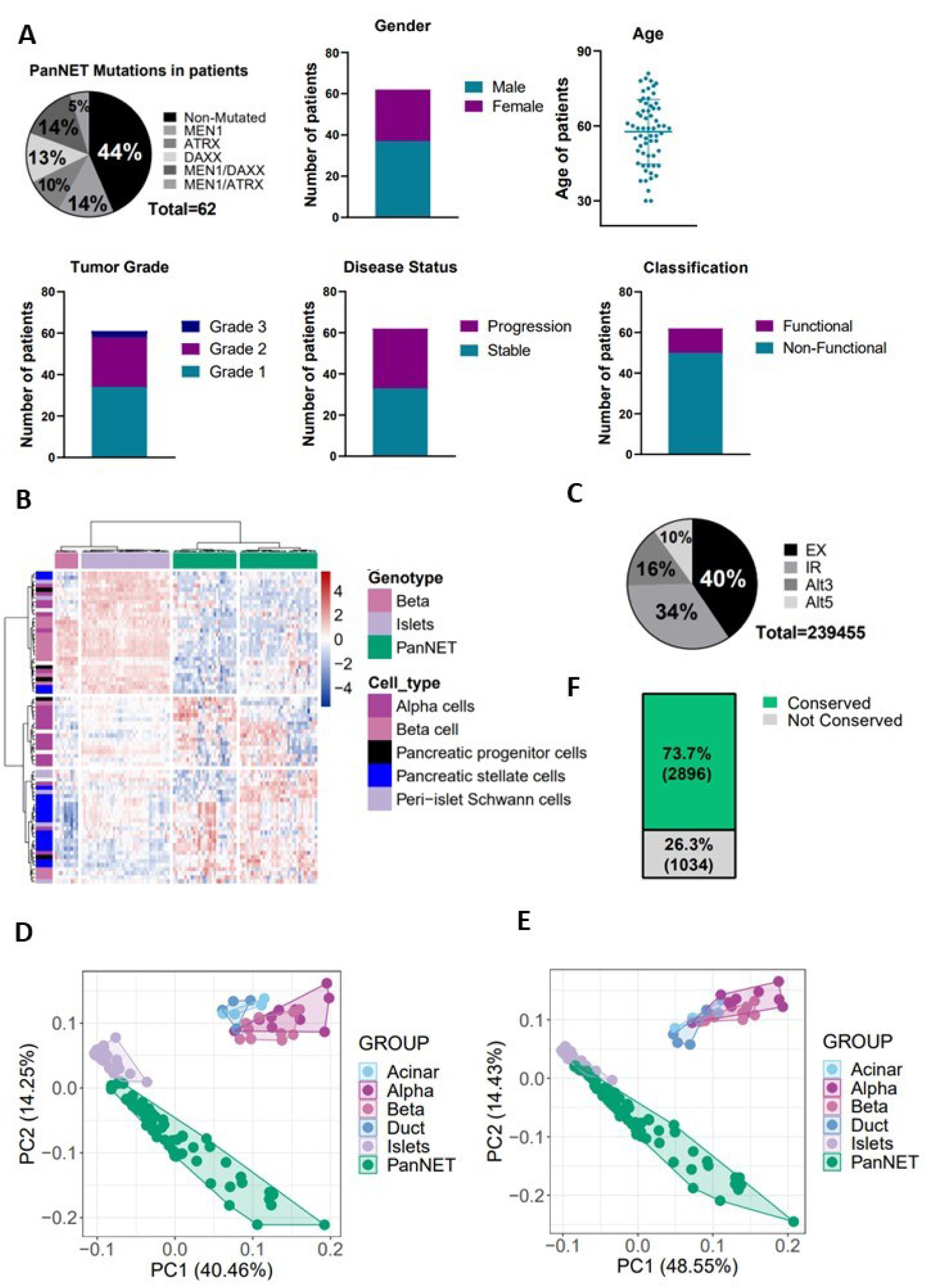
Comparison of alternative splicing in PanNET samples to normal pancreatic tissue reveals the pivotal role of exonic events in neuroendocrine tumors. (**A**) Pie chart depicting the mutational landscape of the tumor samples analyzed, followed by stacked bar graphs and a scatterplot representing the gender, age, tumor grade, disease status and functional classification of the PanNET patients. (**B**) Heat-map depicting the expression profiles of cellular markers of the five pancreatic cell types indicated, with rows representing cell types and columns representing the samples analyzed. (**C**) Pie chart of the distribution of the mapped splicing events in at least 70 out of the 128 samples analyzed in total, based on their type. Color coding shows the different event types (EX: alternative exon, IR: Intron Retention, Alt3: alternative 3’ splice site, Alt5: Alternative 5’ splice site). (**D**), (**E**) PCA analysis of the distribution of tumoral samples in comparison with alpha, beta, acinar, ductal cells or islets, based either on the PSI values of the top 5000 most variable events in general (D) or of the top 5000 most variable intronic splicing events (E). (**F**) Stacked bar graph showing the percentage of the significantly altered spliced events (|ΔPSI|>15, P-Adjusted value<0.05) of our analysis that are conserved between human (hg38) and mouse (mm10) genomes. In brackets, the respective numbers of events.

**Figure 1.**
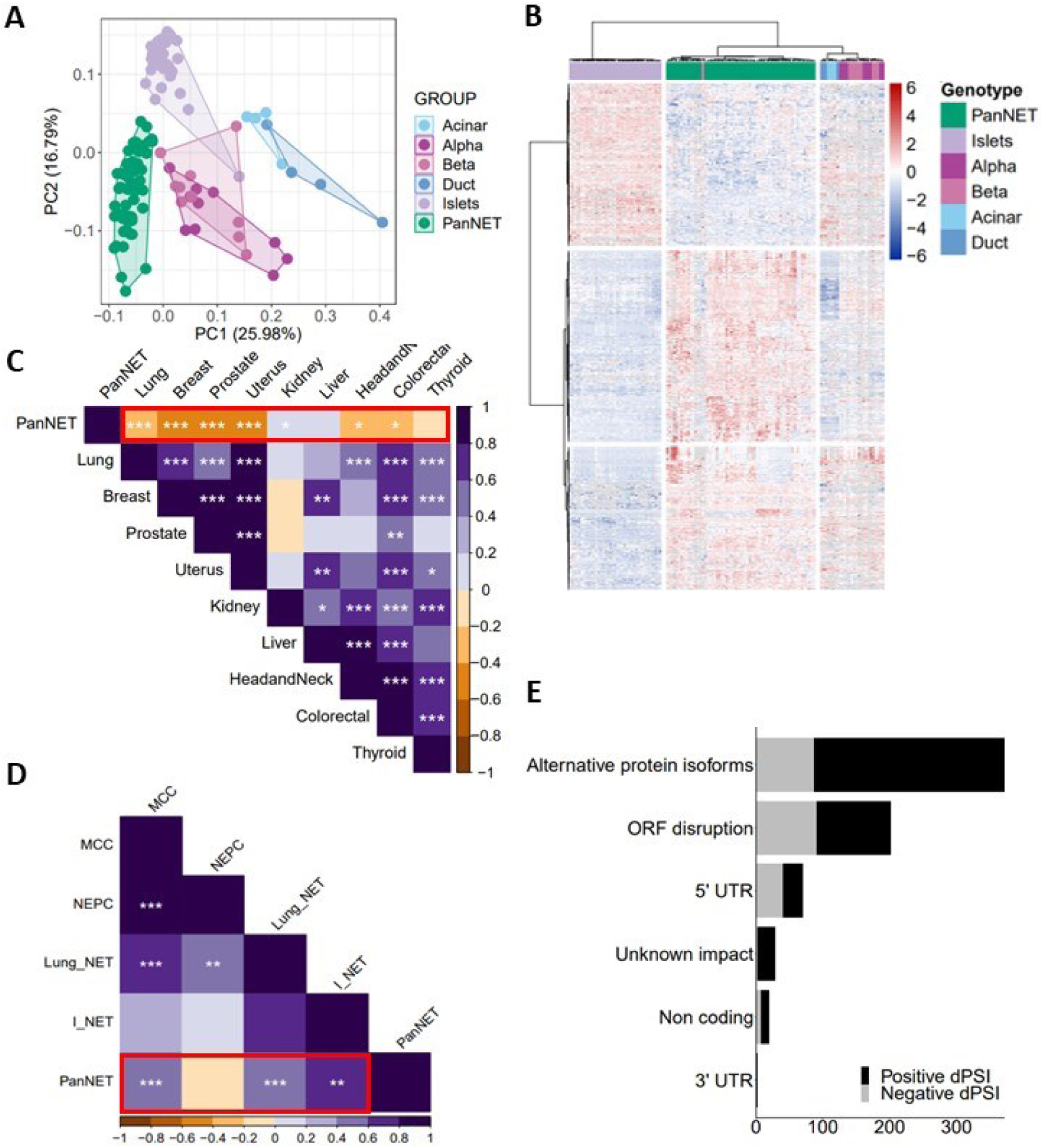
Comparison of alternative splicing in PanNET samples to normal pancreatic tissue reveals the pivotal role of exonic events in neuroendocrine tumors. (**A**) The Principal Component Analysis (PCA) illustrates the distribution of the PSI values of the top 5000 most variable exonic splicing events in tumor samples in comparison with alpha, beta, acinar, ductal cells or islets. Each point represents an individual sample analyzed. (**B**) Heat-map of the top 5000 most variable exonic splicing events showing the clustering of the samples based on their PSI values, with rows representing the events and columns the samples analyzed. (**C**), (**D**) Correlation plots of the common altered exonic events between PanNETs (|ΔPSI|>15, P-Adjusted value <0.05) and other cancers from TCGA database (**C**) or other types of NETs (|ΔPSI|>5, P value <0.05) (**D**). Association values determined by Pearson coefficient are visualized using color grading. Stars represent p-values for the statistical significance of the association value. *,p<0.05; **,p<0.01; ***,p<0.001. (**E**) Stacked bar chart showing the predicted impact that the significantly altered exonic events have on the protein level. The x-axis shows the number of changing events, whereas the y-axis denotes the change caused at the protein level.

Given that there is still great uncertainty as to the cells of origin of PanNETs, we focused mainly on the comparison of alternative exon events between PanNET and normal islets to account for any contribution of all endocrine cell types of the pancreas. Thus, to investigate the magnitude of the differences in inclusion of the alternative exons, we compared their PSI values in PanNETs with the values in normal islets. Upon filtering for the most significantly changed AS events (|ΔPSI| ≥ 15 % and *p*_adjusted_value < 0.05), a total of 704 alternative exon events were identified (Table S1). These events appear to be largely conserved between human and mouse species (73.7% conservation, Figure S1F). The PSI values of the 704 alternative exon events that are significantly different between PanNET and normal islet samples were used to perform an unbiased hierarchical clustering analysis using Euclidean distance (Figure 1B). All the samples were classified in 3 distinct groups, the tumour, islet, and alpha/beta/ductal/acinar group, suggesting that the events under study allow differentiation of the distinct sample types. Furthermore, the larger group of changed events consists of events upregulated (positive ΔPSI) in PanNETs compared to normal islets. Correlation analysis was conducted using the Pearson correlation coefficient to evaluate the association between PanNETs ((|ΔPSI| > 15, P_adjusted value < 0.05, 704 exonic events) and other cancer types from the Cancer Genome Atlas (TCGA) (|ΔPSI| > 5, P value < 0.05) (Figure 1C). PanNETs exhibited negative coefficients, indicating a negative linear relationship with all other cancers. This observation supports our hypothesis that AS changes in PanNETs contribute to a distinctive tumoral phenotype (Figure 1C). Similar correlation analysis between PanNETs (704 altered spliced exonic events) and other NETs (|ΔPSI| > 5, P value < 0.05), revealed the closest similarity of PanNET splicing isoforms with the Intestinal NET ones (I-NET; Figure 1D).

Furthermore, we investigated the impact the changes in the inclusion pattern of the alternative exons have on the respective protein isoforms. As shown in Figure 1D, the majority of these events (∼54%) are predicted to maintain the ORF and thus alter the expression of alternative protein isoforms, while ∼30% of the alternative exons are predicted to disrupt the ORF and affect the final protein levels (Figure 1E).

### The neuronal factor SRRM3 is upregulated in PanNETs and drives the deregulation of an islet splicing program

To identify splicing regulators possibly mediating the observed AS changes we examined the global gene expression data of the PanNET samples for potential changes in the levels of splicing factor mRNAs in comparison to the normal islet datasets. A stringent analysis (|Log_2_FoldChange|>2 and P_adjusted value<0.05) led to the identification of 23 splicing factors significantly upregulated in PanNETs compared to normal islet samples (Table S2). Of these, 7 are known to regulate neuronal splicing (Figure 2A). To identify which of the significantly deregulated splicing factors can be responsible for the AS changes noted, we performed a splicing Factor-Event Interface Analysis, which scrutinized the interplay between differentially expressed splicing factors (|log_2_FoldChange|>0.58, P_adjusted value<0.5) and changes in AS events in PanNETs. This analysis clearly suggested that the neuronal, regulatory splicing factor SRRM3 controls a substantial portion of the detected AS changes (Figure 2B, C). The majority of these exons are positively regulated (more included in the final mRNA product) in PanNETs, where SRRM3 is upregulated. Furthermore, investigating the presence of the known SRRM3 binding motif (UGC) in the sequence of the deregulated alternative exons and their adjacent introns showed the expected position of the SRRM3 binding motif upstream from the majority of the upregulated exons^28^ (Figure S2A).

**Figure 2.**
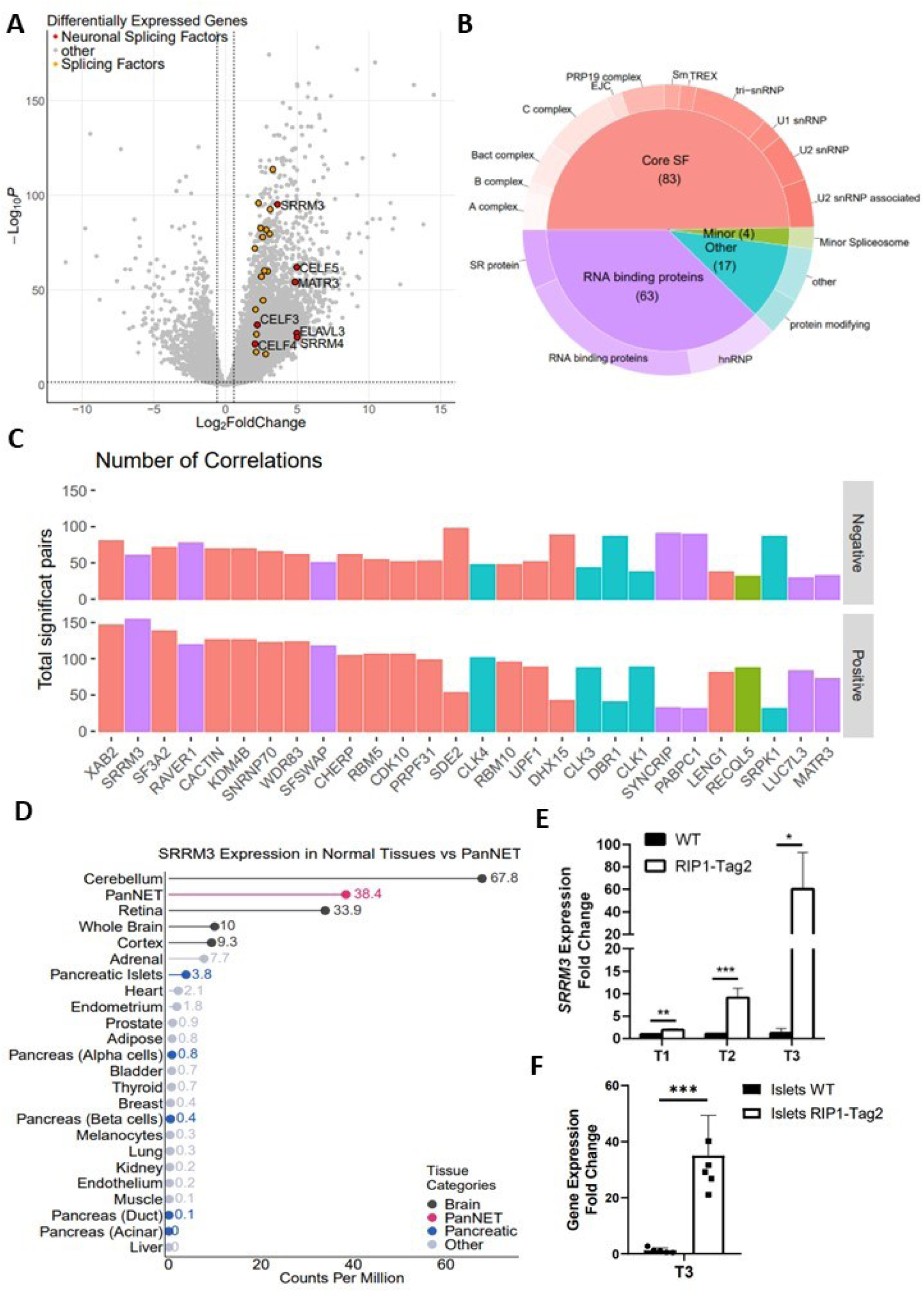
The neuronal splicing factor SRRM3 is significantly upregulated and controls splicing deregulation in PanNETs. (**A**) Volcano plot of the log_2_FoldChange values in mRNA levels between PanNET samples and normal pancreatic islets. In color, 23 splicing factors were found significantly upregulated in PanNET samples (|Log2FoldChange|>2, P-Adjusted value<0.05), 7 of which are known to regulate neuronal splicing (depicted in red and named). (**B**) Pie-Donut chart of the splicing factors that were found deregulated in the differential expression analysis (|Log_2_FoldChange|>0.58, P-Adjusted value<0.05), organized in classes based on splicing-related proteins. Colors depict different categories as indicated. (**C**) Bar-plot illustrating the number of correlations (positive and negative) of the deregulated splicing factors from panel B with the significantly altered exonic events from the splicing analysis (|ΔPSI|>15, P-Adjusted value<0.05), in a descending order. Colors depict different splicing classes as shown in panel B. (**D**) Lollipop chart showing SRRM3 average cRPKM values from the differential expression analysis regarding PanNET and normal pancreatic tissues analyzed in comparison to other normal tissues. Data acquired from VastDB^15^. Color coding corresponds to organized categories as indicated. (**E**) Bar-plot of RT-qPCR analysis of SRRM3 average expression in WT and RIP1-Tag2 mice in three different timepoints (T1, T2, T3). GAPDH levels were used for the normalization. Two-tailed unpaired t-test was used to detect statistically significant changes. *, p<0.05; **, p<0.01; ***, p<0.001. (**F**) Bar-plot of RT-qPCR analysis of SRRM3 expression in islets isolated from WT and RIP1-Tag2 mice of T3 timepoint. GAPDH levels were used for the normalization. Two-tailed unpaired t-test was used to detect statistically significant changes. ***, p<0.001.

**Figure S2.**
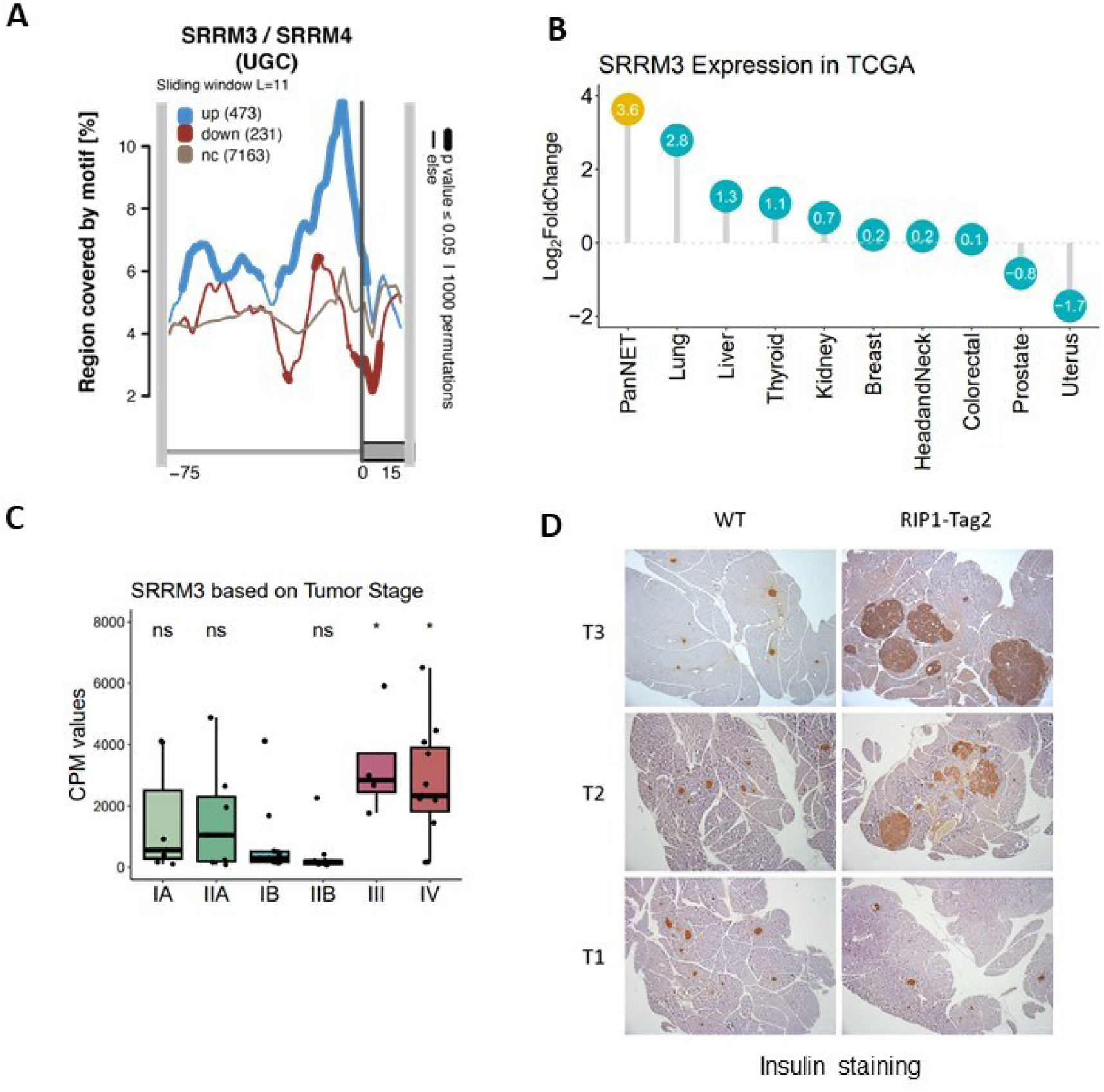
The neuronal splicing factor SRRM3 is significantly upregulated and controls splicing deregulation in PanNETs. (**A**) RNA map representing the distribution of SRRM3/SRRM4 binding motif in alternative regulated exons (down: decreased inclusion; up: increased inclusion; nc: non-changing between PanNET and normal islet exons) and upstream introns, compared to control. Thicker segments indicate regions in which enrichment is significantly different over non-changing (p-value<0.05). The number of events in each category are shown in brackets. (**B**) Lollipop chart showing SRRM3 Log_2_FoldChange values in PanNETs compared to other types of cancer from the TCGA repository. Differential expression values are as shown, and colors distinguish the highest expression value of SRRM3 in PanNETs. (**C**) Box plot of the averages of counts per million (CPM) values for SRRM3 in PanNET patients categorized according to their tumor stage. Each point represents an individual sample analyzed. Individuals with no clinical information about tumor stage were not included in this graph. *P* values were calculated using a one-way ANOVA test. ***,p<0.001; ****,p<0.0001. (**D**) WT and RIP1-Tag2 pancreases from all timepoints of the disease were embedded in paraffin, sectioned and stained for Hematoxylin and also probed with antibodies against Insulin. Representative pictures from every timepoint are displayed, showing increased insulin levels in RIP1-Tag2 tissues.

SRRM3 is a splicing factor enriched in neuronal tissues and particularly in brain cerebellum, whereas the expression levels of the SRRM3 mRNA in pancreas are very low. In PanNETs *SRRM3* mRNA levels increase dramatically, reaching neuronal tissue levels (Figure 2D). Comparative analysis of the deregulation of the *SRRM3* mRNA levels in PanNETs and in different TCGA cancers revealed that in PanNETs SRRM3 exhibits the highest levels of deregulation (Figure S2B). Furthermore, *SRRM3* levels are higher in the later PanNET stages, III and IV according to the patients’ data that we analysed (Figure S2C).

We used the RIP1-Tag2 (RT2) mouse model^29^ to assay the levels of SRRM3 in PanNETs. PanNETs occur in RT2 mice due to expression of the SV40 T-antigen oncoprotein (Tag) from a rat insulin promoter (RIP). Tumour formation in RT2 mice is rapid and synchronized. Tumours in this model arise mainly in beta-cells and are thus insulinomas. We assayed the expression levels of SRRM3 in pancreatic RNA isolated from RT2 mice at three distinct time points: T1=8w, T2=12w and T3=16-17w. The three time points of the RIP1-Tag2 mice that are on a high glucose diet were selected based on the observations that RT2 mice at Τ1 show T-antigen expression in pancreatic islets, but no signs of islet hyperplasia. At Τ2, islet hyperplasia can be noticed, whereas at Τ3 growth of large, insulin producing tumours is easily detected^29,30^ (Figure S2D). Using qPCR, we tested *SRRM3* mRNA expression in the pancreas of RT2 and WT mice at the three time points (Figure 2E). Interestingly, *SRRM3* mRNA had a two-fold increase in the pancreas of RT2 mice already at the 1st time point and the overexpression became more and more apparent with tumour progression, reaching a 60-fold increase in the pancreas of mice at the last time point. *SRRM3* mRNA overexpression was also apparent in islets isolated from the pancreas of RT2 mice when compared to islets from WT mice (Figure 2F).

### SRRM3 participates in neuroendocrine cell dysfunction via regulation of AS of a group of neural microexons

It is known that SRRM3 controls the splicing of microexons smaller than 27nts^28^, particularly in locations where SRRM4 is not expressed^31^, which is the case for a small subset of the PanNETs analysed (Figure S3A). Recently, SRRM3 was reported as an important regulator for glucose homeostasis in pancreatic islets, via regulation of AS of a conserved subset of microexons^32^ in pancreatic beta cells. Intriguingly, our analysis reveals that the vast majority of the altered microexons (83%) seem to be upregulated in PanNETs compared to normal islets, with a broader range of ΔPSI values, compared to other upregulated exonic events, thus pinpointing to a significant role of microexons in PanNET regulation (Figure 3A). Comparative analysis of the microexon events identified by Juan-Mateu et al^32^ as deregulated upon *SRRM3* knock-down in the pancreatic beta-cell line EndoC-betaH1, with the 179 microexons identified as deregulated in PanNETs by our analysis, revealed a very weak negative correlation between the two microexon groups (Figure S3B) with 51 common events (Figure S3C). A positive correlation was observed, when we compared the 179 microexons that are deregulated in PanNETs with the group of neural (N) and pancreatic/neural (PN) microexons that are deregulated in HeLa cells upon overexpression of *SRRM3* (Figure 3B)^32^. Furthermore, the majority of the mRNAs affected by the differential inclusion of microexons in PanNETs are genes that bear microexons characterized as important for neurogenesis^19^ (Figure 3C).

**Figure 3.**
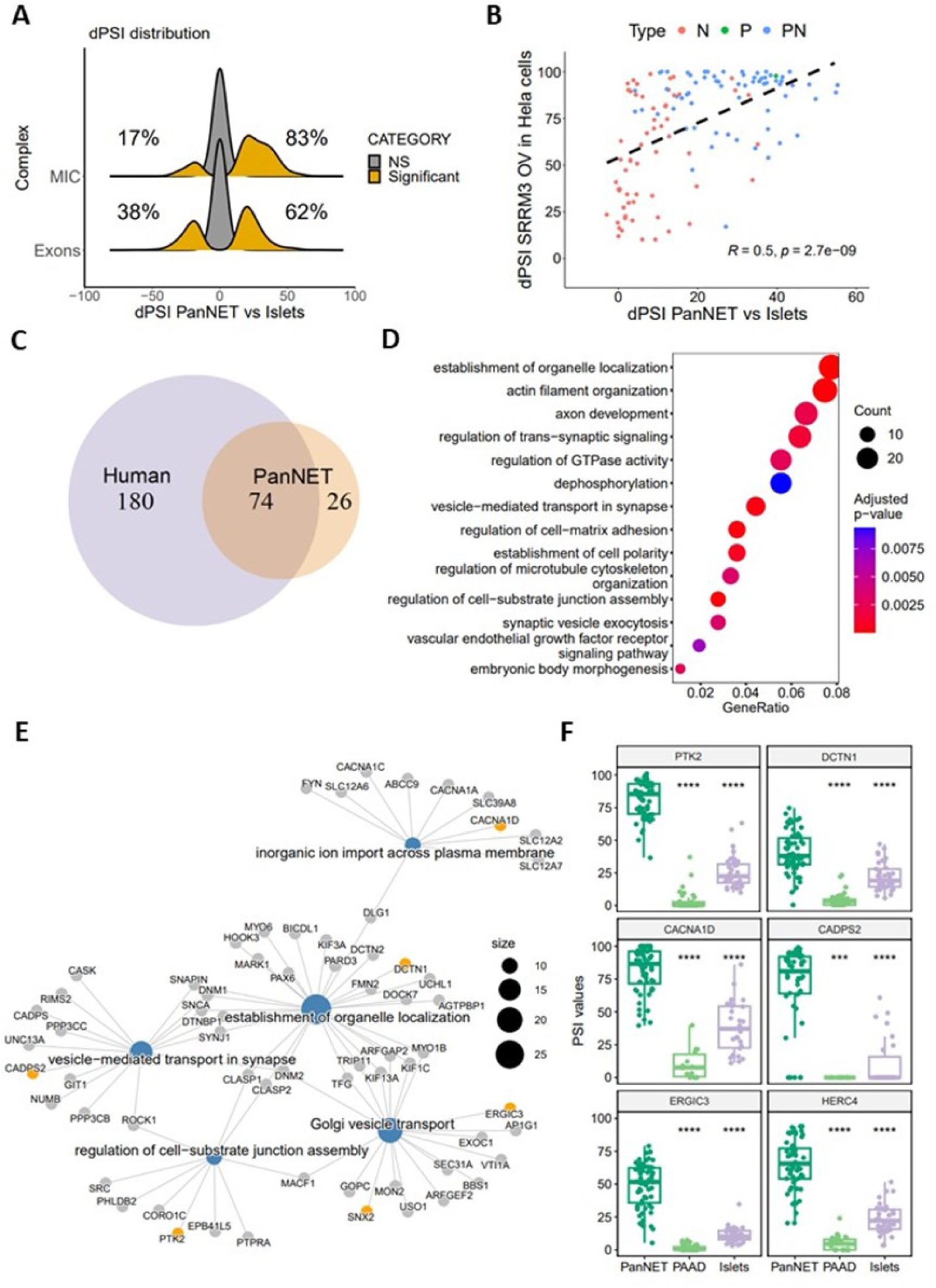
Characterization of a group of highly upregulated microexons in PanNETs compared to normal pancreatic islets. (**A**) Plot showing the distribution of the ΔPSI values for exonic and MIC events in PanNETs compared to normal islets. Non-significant (NS) events are presented in grey and in orange are the significantly changing ones. Percentages represent the distribution of ΔPSI values of the events being either positive or negative. (**B**) Scatterplot of the SRRM3-regulated MIC events that are common between the list of upregulated microexons in HeLa cells overexpressing *SRRM3*^32^ and our analysis. A positive correlation is calculated (R = 0.5, P value = 2.7 x 10^−09^). N: neural, P: pancreatic, PN: pancreatic and neural microexons. (**C**) Venn-diagram of genes with deregulated microexons in PanNETs and genes with microexons characterized as neuronal in human^19^. (**D**) Dot plot depicting the result from the Gene Ontology-Biological Process (GO-BP) enrichment analysis after further reduction using the Revigo tool^33^. Only microexon events (Length <50) with P-Adjusted value<0.05 were used. Color coding indicates the adjusted p-value range, while the size depicts the number of genes involved in each process as indicated. (**E**) Gene-concept network (cnetplot) of genes across highly enriched GO BP in PanNETs compared to islets. Color coding reflects the genes of interest. The size of the BP nodes reflects the number of genes with altered AS that belong in this BP. (**F**) Box plots of the PSI values for each selected splicing event. Each point represents an individual sample analyzed. P values were calculated using a one-way ANOVA test. ***,p<0.001; ****,p<0.0001. PAAD: Pancreatic adenocarcinoma.

**Figure S3.**
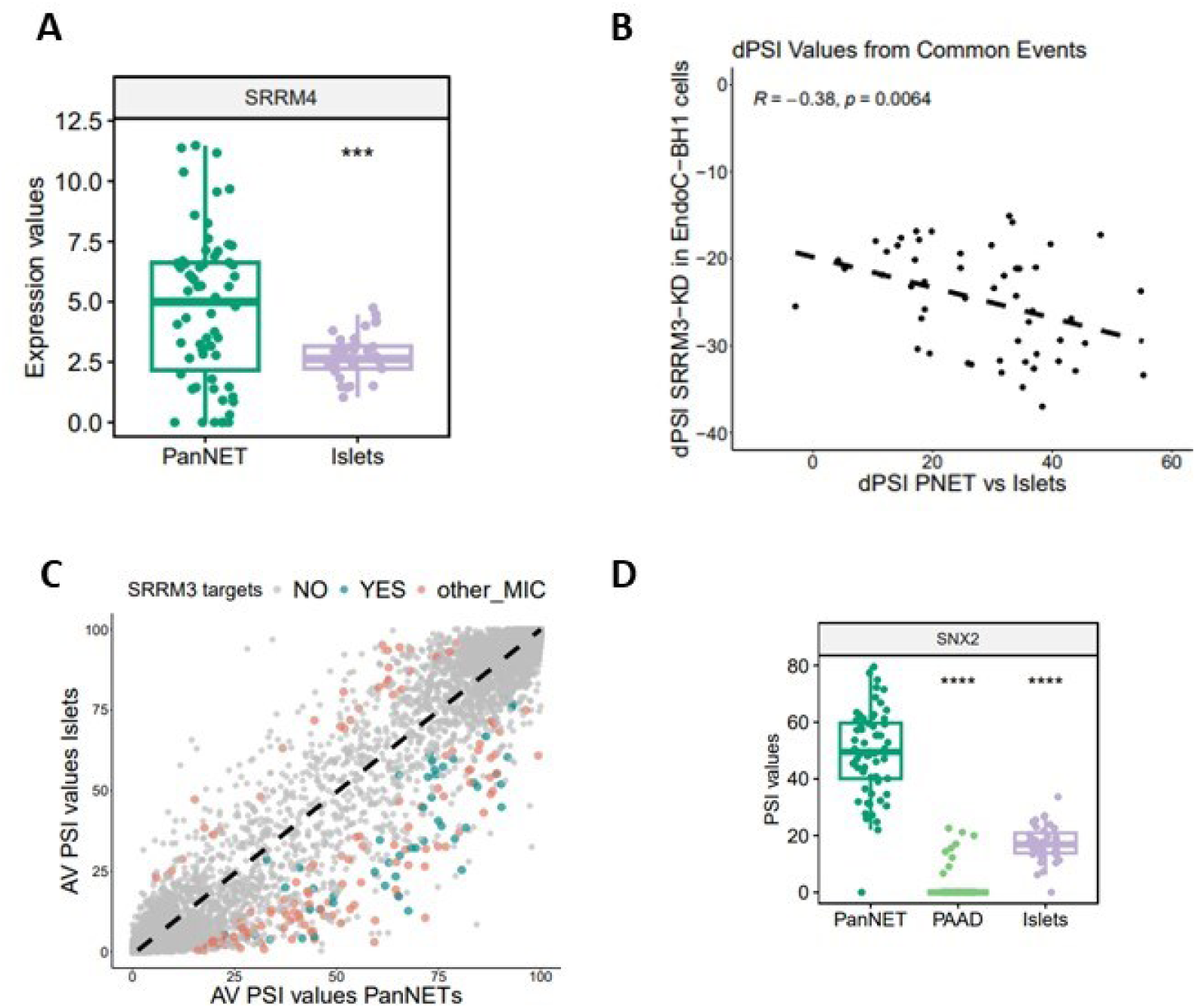
Characterization of a group of highly upregulated microexons in PanNETs compared to normal pancreatic islets. (**A**) Box plot of the log2 expression values for *SRRM4* mRNA in PanNET and normal islets. Each point represents an individual sample analyzed. *SRRM4* expression was not detected in PAAD samples. P values were calculated using an one-way ANOVA test. ***, p<0.001. (**B**) Scatterplot of the 51 SRRM3-regulated microexon events that are common between the list of downregulated microexons in *SRRM3*-Knhock-down EndoC-betaH1 cells^32^ and our analysis. A moderate, negative correlation is calculated (R = –0.38, P value = 0.0064). (**C**) Scatterplot of the average PSI value in PanNETs compared to normal islets. In cyan color, microexons (P-Adjusted value <0.05 in PanNETs compared to islets) known to be direct targets of SRRM3^32^. In orange, the rest of significantly altered MIC from our analysis. (**D**) Box plot of the PSI values for the SNX2 MIC, the only selected MIC from our analysis that is not conserved between human and mouse genome. Each point represents an individual sample analyzed. P values were calculated using a one-way ANOVA test. ****,p<0.0001.

To investigate the biological roles of the deregulated microexons in PanNETs, we analysed the biological processes (BPs) the affected genes are associated with. We used Revigo^33^ to remove redundant GO terms, after which axon development, vesicle mediated transport, synaptic vesicles and signalling were amongst the most enriched BPs, pinpointing to the deregulation of a neuronal phenotype in PanNETs (Figures 3D). Given that these GO terms signify cellular functions that are critical for neuroendocrine tissue homeostasis, which is impaired or problematic in PanNETs, we focused our analyses on the impact AS has in such processes. For this, we specifically pursued the investigation of a representative group of alternatively included microexons that depict high ΔPSI values between PanNET and normal islet samples and are associated with these categories: CADPS2 (Calcium-dependent secretion activator 2), PTK2/FAK1 (Protein-tyrosine kinase 2/Focal adhesion kinase 1), CACNA1D (Voltage-dependent L-type calcium channel subunit alpha-1D), HERC4 (HECT domain and RCC1-like domain-containing protein 4), ERGIC3 (Endoplasmic reticulum-Golgi intermediate compartment protein 3), DCTN1 (Dynactin subunit 1), SNX2 (Sorting nexin-2) (Table S1 and Figure 3E). According to our analyses, the respective alternative microexons are highly included in PanNETs compared to normal islets or to other types of pancreatic cancers (Figure 3F and S3D). Importantly, with the exception of SNX2, the selected events are conserved between humans and mice.

To validate the change in the inclusion percentage for the above mentioned alternative microexons during PanNET progression, we used RNA from the pancreas of the RT2 mouse model, at the three time points described above. For all the microexons tested we could detect increased inclusion already at T2 of the RT2 mice, compared to their littermate wt mice and even more profound increase at T3 (Figure 4A) following the overexpression pattern of SRRM3 (Figure 2E). For certain cases, the increased inclusion in RT2 vs WT was obvious even at T1 (Figure 4A, *Cacna1D*, *Ptk2*). Analysis of RNA from individual tumours agreed to the elevated inclusion levels of the microexons in RT2 mice (Figure 4B). Use of RNAs from mouse pancreatic islets allowed validation of the increased inclusion levels of the microexons in RT2 mice compared to their littermate wt mice, however, they depicted smaller inclusion percentages (Figure S4A). Increased insulin and glucagon levels in the isolated islets verified their successful isolation (Figure S4B), however the reduced inclusion percentages could possibly be explained by the fact that islets derived from tumours are a mixture of normal, and hyperplastic islets^34^.

**Figure 4.**
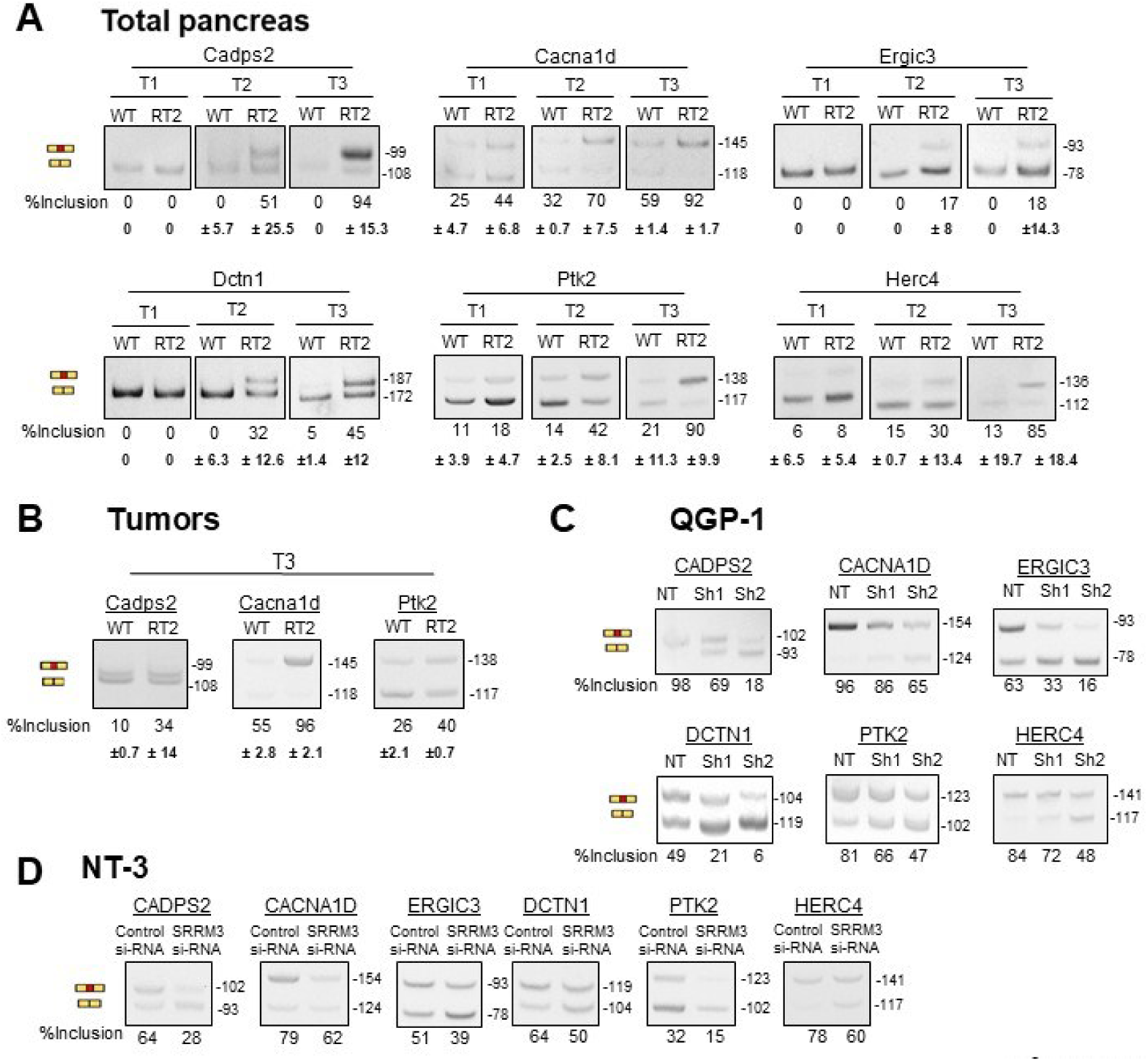
SRRM3 regulates the alternative splicing of neural microexons during PanNET progression. (**A**) Analysis by RT-PCR and gel electrophoresis of the indicated AS events events (*Cadps2*, *Cacna1d*, *Ergic3*, *Dctn1*, *Ptk2*, *Herc4*) during PanNET progression, using RNAs isolated from total pancreatic tissues of WT and RT2 (RIP1-Tag2) mice, at the time points indicated (T1, T2, T3). The %inclusion values represent the quantification of the inclusion band relative to the total signal from both the inclusion and skipping bands. Error bars, based on the standard deviation (±SD) values from the corresponding biological replicates, are depicted below. Product lengths (bp) are marked on the right of each picture. (**B**) Similar RT-PCR analysis of the indicated events in RNA samples generated from isolated tumors from RT2 mice, at the timepoint T3. (**C**) RT-PCR analysis of the indicated events in RNA samples derived from QGP-1 control non-treated cells (NT) and *SRRM3* knock-down QGP1 cells generated using two different *SRRM3* specific shRNAs (sh1, sh2). (**D**) RT-PCR analysis of the indicated events in RNA samples derived from NT-3 control cells (treated with scramble siRNAs) or NT-3 *SRRM3* knock-down cells (treated with *SRRM3* specific siRNAs).

**Figure S4.**
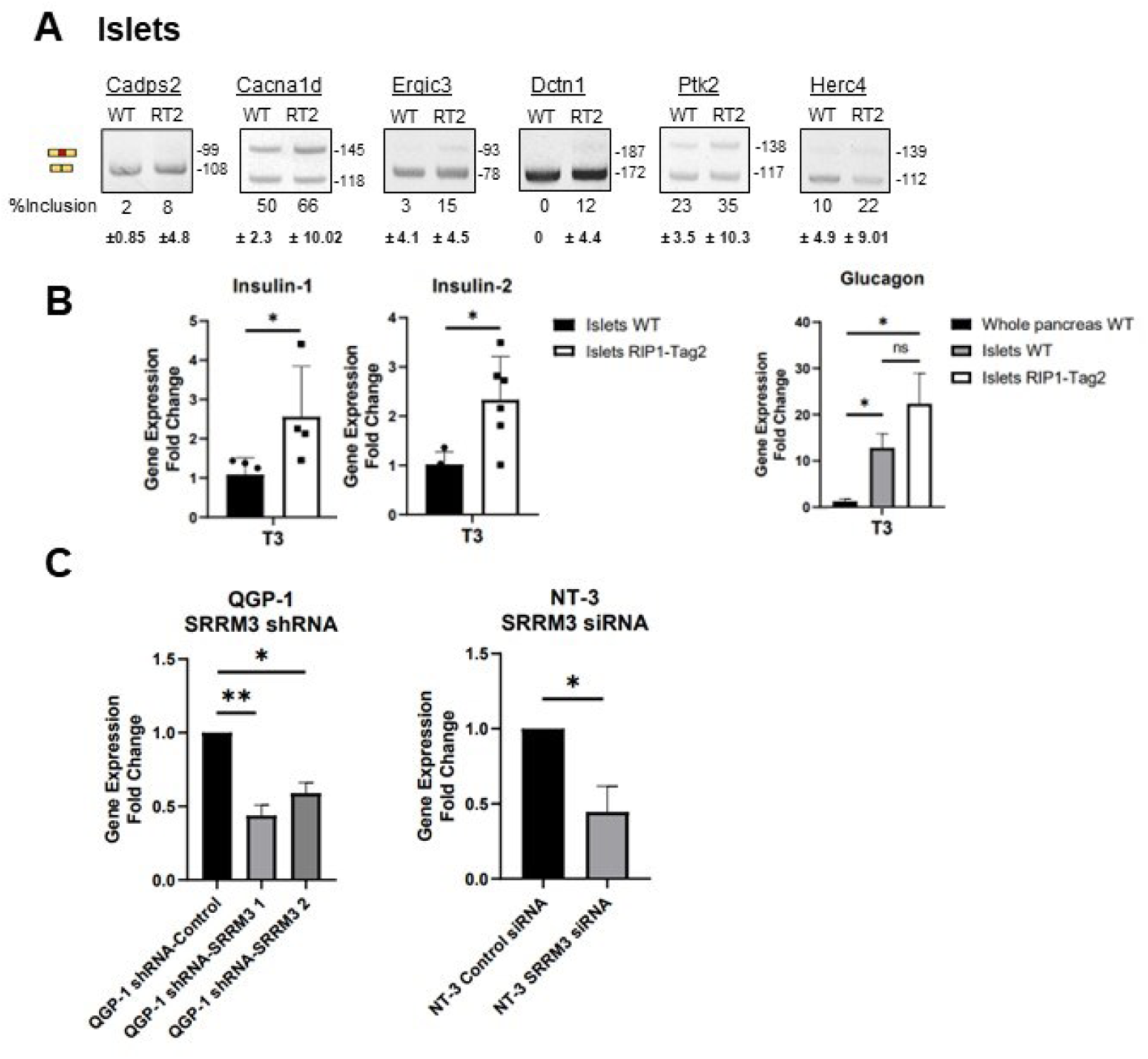
SRRM3 regulates the alternative splicing of neural microexons during PanNET progression. (**A**) Analysis by RT-PCR and gel electrophoresis of the indicated AS events (*Cadps2*, *Cacna1d*, *Ergic3*, *Dctn1*, *Ptk2*, *Herc4*), using RNAs from isolated murine pancreatic islets from WT and RIP1-Tag2 (RT2) mice. The %inclusion values represent the quantification of the inclusion band relative to the total signal from both the inclusion and skipping bands. Error bars, based on the standard deviation (±SD) values from the corresponding biological replicates, are indicated below. Molecular lengths (bp) are marked on the right of each picture. (**B**) Bar-plots illustrating the average expression levels of *Insulin 1*, *Insulin 2*, and *Glucagon* as they were guantified by RT-qPCR using RNA from islets of WT and RIP1-Tag2 (RT2) mice. Comparison to the whole pancreatic levels is shown for *Glucaton* at timepoint T3. *GAPDH* levels were used for the normalization in *Insulin 1* and *Insulin 2*. HPRT levels were used for the normalization in *Glucagon* instead of *GAPDH*. Two-tailed unpaired t-test was used to detect statistically significant changes. *, p<0.05. (**C**) Bar-plots of RT-qPCR analyses of SRRM3-Knock-down effectiveness in QGP-1 and NT-3 cells. GAPDH levels were used for the normalization. Two-tailed unpaired t-test was used to detect statistically significant changes. *, p<0.05; **, p<0.01. (**D**) RT-qPCR analysis of the overexpression levels of *SRRM3* in EndoC-βH3 cells, induced by increasing amount of doxycycline (Dox) as indicated. (**E**) Analysis by RT-PCR and gel electrophoresis of the indicated AS events (*CADPS2*, *CACNA1D*, *ERGIC3*, *DCTN1*, *PTK2*, *HERC4*) using RNAs from EndoC-βH3 cells overexpressing *SRRM3* after induction with doxycycline (Dox) at the indicated concentrations. The %inclusion values represent the quantification of the inclusion band relative to the total signal from both the inclusion and skipping bands.

To verify the dependency of these events on SRRM3, we used the PanNET cell lines QGP-1 and NT-3. QGP-1 cells have been derived from a primary pancreatic carcinoma, expressing mainly somatostatin and very low levels of insulin^35^, whereas NT-3 have been derived from the lymph node of a patient with well differentiated NET of the pancreas, expressing high levels of insulin^35^. We downregulated the levels of *SRRM3* mRNA in QGP-1 cells by stably transfecting with constructs that result in expression of two different shRNAs specific for *SRRM3* or a non-specific (NT) shRNA. The different shRNAs for SRRM3 that were used produce varying degrees of silencing of SRRM3 (Figure S4C). For NT-3 cells, the levels of SRRM3 were downregulated using specific siRNAs against *SRRM3* and scrambled siRNAs as a control. In all cases, the effectiveness of the downregulation of *SRRM3* was verified by qPCR (Figure S4C). Upon verification of *SRRM3* downregulation, we performed RT-PCR to assay the inclusion levels of the microexons of interest (Figure 4C, 4D). For both PanNET cell lines we could detect the expected decrease in inclusion of the microexons upon downregulation of *SRRM3*.

### *SRRM3* controls the response of intracellular calcium levels to stimuli in PanNET cells

There is functional evidence for the close association of calcium levels with secretory granule exocytosis in normal pancreatic β cells *in vitro*^36^. Interestingly, the microexons that depict the highest ΔPSI between PanNETs and normal islets and are controlled by SRRM3 affect microdomains of proteins that participate in this highly orchestrated process in islets and neuronal cells. For example, the alternative microexon that is almost exclusively included in the *CADPS2* pre-mRNA in PanNETs (exon 19) is a brain specific neuronal microexon^19^ that adds 3 amino acid residues between two alpha helices in the coiled-coiled domain of the CADPS2 protein (Figure 5A). Importantly, this event has not been detected as altered in other NETs or TCGA cancers (Tables S3, S4). CADPS2 is of fundamental importance for priming synaptic vesicles for fusion and neurotransmitter release in neurons, for Large Dense Core Vesicle (LDCV) secretion and for the stability and recruitment of insulin granules in mouse pancreatic beta cells^37,38^. The protein domain that is altered by the included microexon is responsible for interaction with the D2 dopamine receptor, with no other known functions so far^39^. The significantly high inclusion of the alternative microexon of *CACNA1D* pre-mRNA (exon 44, which is also a brain specific microexon^19^), adds 9 amino acids at the C-terminal cytoplasmic region of the L-Type Calcium channel CaV1.3 (Figure 5A), a protein important for glucose-induced insulin secretion by pancreatic islets^40^. Inclusion of the microexon results in the expression of a protein isoform with reduced calcium dependent inactivation capability in mouse brain^41^. The inclusion of a microexon in *PTK2* pre-mRNA (exon 6; Figure 5A) leads to the expression of a FAK1 protein isoform with altered levels of glucose induced phosphorylation of Tyr397 and this change affects the docking of insulin granules at plasma membrane of pancreatic beta cells^42,43^. Similarly, the microexons included predominantly in PanNETs in the pre-mRNAs of *ERGIC3*, *DCTN1*, *HERC4* are almost exclusively brain specific^19^ and alter the respective protein isoforms, with unknown functional outcome to date (Figure S5A). The above-mentioned isoforms have not been connected yet to neuroendocrine tumour development and progression.

**Figure 5.**
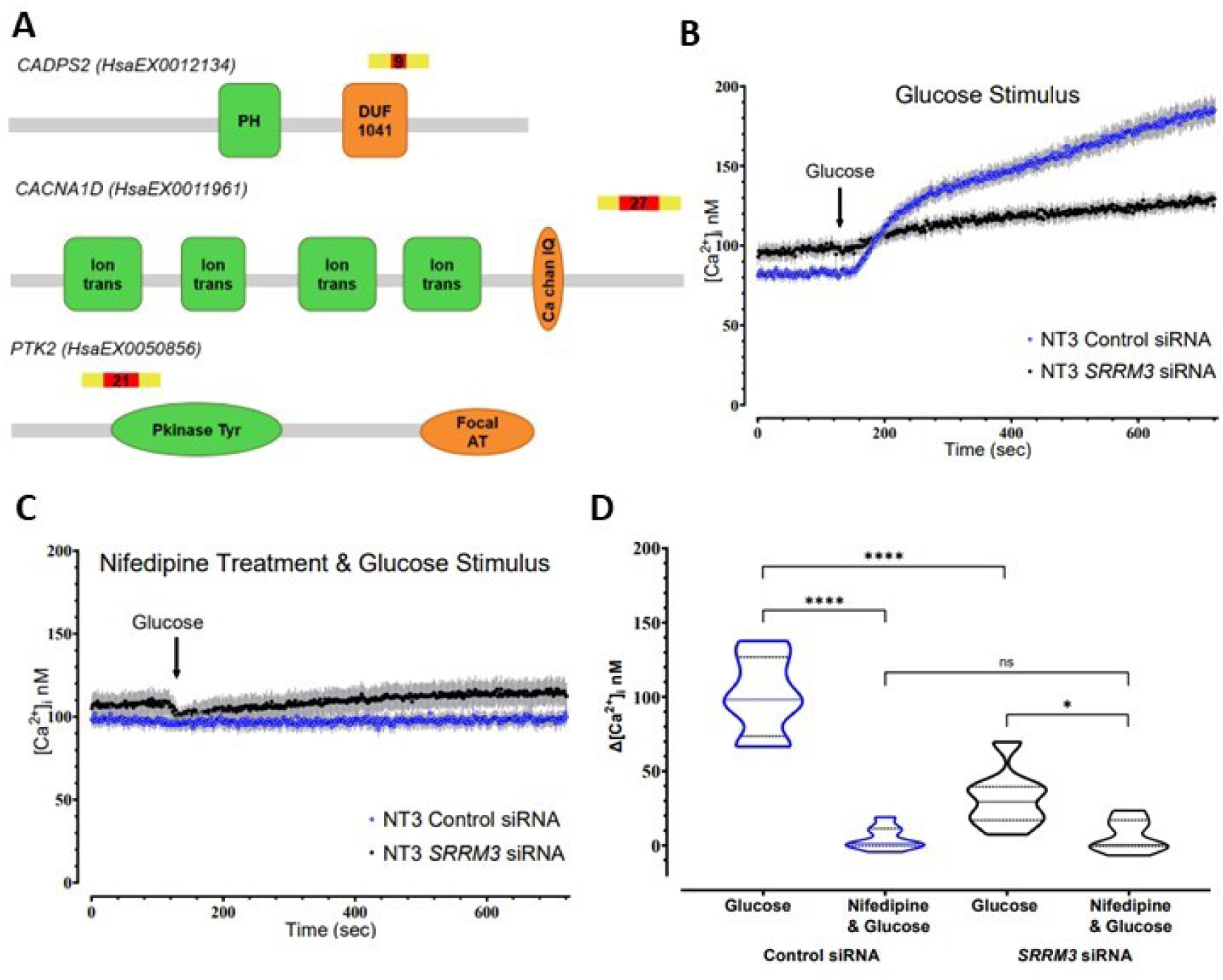
SRRM3 regulates intracellular calcium responses in PanNET cell lines. (**A**) Schematic showing the anticipated effect of alternative splicing events on the resulting proteins CADPS2, CACNA1D, PTK2. The position of the alternative exons in relation to the known functional domains are marked in each case. PH: Pleckstrin Homology, DUF1041: Domain of Unknown Function 1041, Ion trans: Ion transport, Ca Chan IQ: IQ motif within the Ca²⁺ channel, Pkinase Tyr: Protein Tyrosine Kinase, Focal AT: Focal Adhesion Targeting domain. (**B**) (**C**) [Ca^2+^]_i_ levels of NT-3 scramble siRNA control or SRRM3 siRNA cells. Error bars represent ±SEM. The arrows indicates the timepoint (120 sec) at which the stimulus (20mM Glucose) was added. In (**C**) nifedipine (10μM) was added in the wells 5 min prior to addition of Glucose. (**D**) Violin plots summarize the differences in [Ca^2+^]_I_ (Δ [Ca^2+^]_i_) at the start and end of intracellular calcium measurements shown in B and C. Two-tailed unpaired t-tests were used to detect statistically significant changes.*, p<0.05; ***, p<0.001.

*SRRM3* was recently shown to be necessary for proper regulation of insulin secretion in non-tumoral mouse islets and required for normal islet architecture^32^. However, its role in PanNET development has not been studied. The fact that SRRM3 controls the inclusion of microexons that alter the functionality of proteins in secretory vesicle trafficking and exocytosis prompted us to assess the involvement of SRRM3 on the secretory ability of PanNET cells in response to stimuli. For this and given that Cav1.3 is an L-type voltage gated calcium channel, highly expressed in pancreatic cells and CADPS2 binds to Ca^2+^ to function in secretory vesicle exocytosis, we firstly tested whether SRRM3 is critical for changes in intracellular calcium levels upon stimulation by glucose or KCl in PanNET cell lines *in vitro*.

More specifically, NT-3 cells transfected with siRNAs specific for SRRM3 (*SRRM3* siRNA) or with scrambled-siRNA (control siRNA) (Figure S4C) were incubated with 20mM glucose and the intracellular calcium levels were assayed using Fluo-4/AM. Control cells responded to the presence of glucose by exhibiting a robust increase in [Ca^2+^]i (Figure 5B). In the presence of 10 μΜ nifedipine, an inhibitor of L-type voltage gated calcium channels that was added to the cells 5 min before addition of glucose, the increase in [Ca^2+^]i was eliminated (Figure 5C). NT-3 cells with reduced *SRRM3* levels showed a substantially attenuated response to the glucose stimulus (Figure 5B). The small response of NT3-KD cells to glucose, probably attributed to the non-complete downregulation of *SRRM3* in these cells, was eliminated by the addition of 10 μΜ nifedipine which once again completely abolished the response of these cells to 20 mM Glucose (Figure 5C). By plotting the responses of the NT3 scrambled-siRNA control cells and *SRRM3* downregulated cells (Figure 5D) it became obvious that a decrease in *SRRM3* expression leads to much smaller ability of these PanNET cells to increase intracellular calcium in response to glucose. This increase in intracellular calcium levels is mediated through L-type voltage gated Ca^2+^ channels since it is diminished in the present of nifedipine (Figure 5D).

In a slightly different approach, QGP-1 cells with reduced expression of *SRRM3* and control cells (Figure S4C) were also assayed for [Ca^2+^]i using KCl to depolarize the cells, because these cells do not respond greatly to stimulation by glucose^44^. Addition of 50 mM KCl in the cells resulted in a sharp and robust spike in [Ca^2+^]i for all 3 cell lines before [Ca^2+^]_i_ levelling off (Figure S5B). The highest increase was recorded in the QGP-1 cells treated with non-specific shRNAs. A close inspection of the observed spikes in [Ca^2+^]i (Figure S5B) revealed that in QGP-1 cells the increase in [Ca^2+^]i reached its peak at 11 sec after addition of KCl, whereas in QGP-1-sh1 cells the peak in [Ca^2+^]i was significantly smaller and peaked at 7 sec and in QGP-1-sh2 the peak was also smaller and peaked at 9 sec after addition of KCl (Figure S5B). Once again, the robust spikes in [Ca^2+^]i generated by KCl were eliminated in all three cell populations in the presence of 10 μΜ nifedipine (Figure S5C, D). Interestingly, blockage of Cav1.3 in PanNET cells, results in reduced viability, as was detected by MTT assays in QGP-1 cells treated with nifedipine (Figure S5E). Together these results suggest that in PanNET cells, SRRM3 promotes the formation of a splicing variant of the voltage-gated Ca^2+^ channel Cav1.3, that exhibits altered response kinetics upon cellular stimulation and supports cell viability.

**Figure S5.**
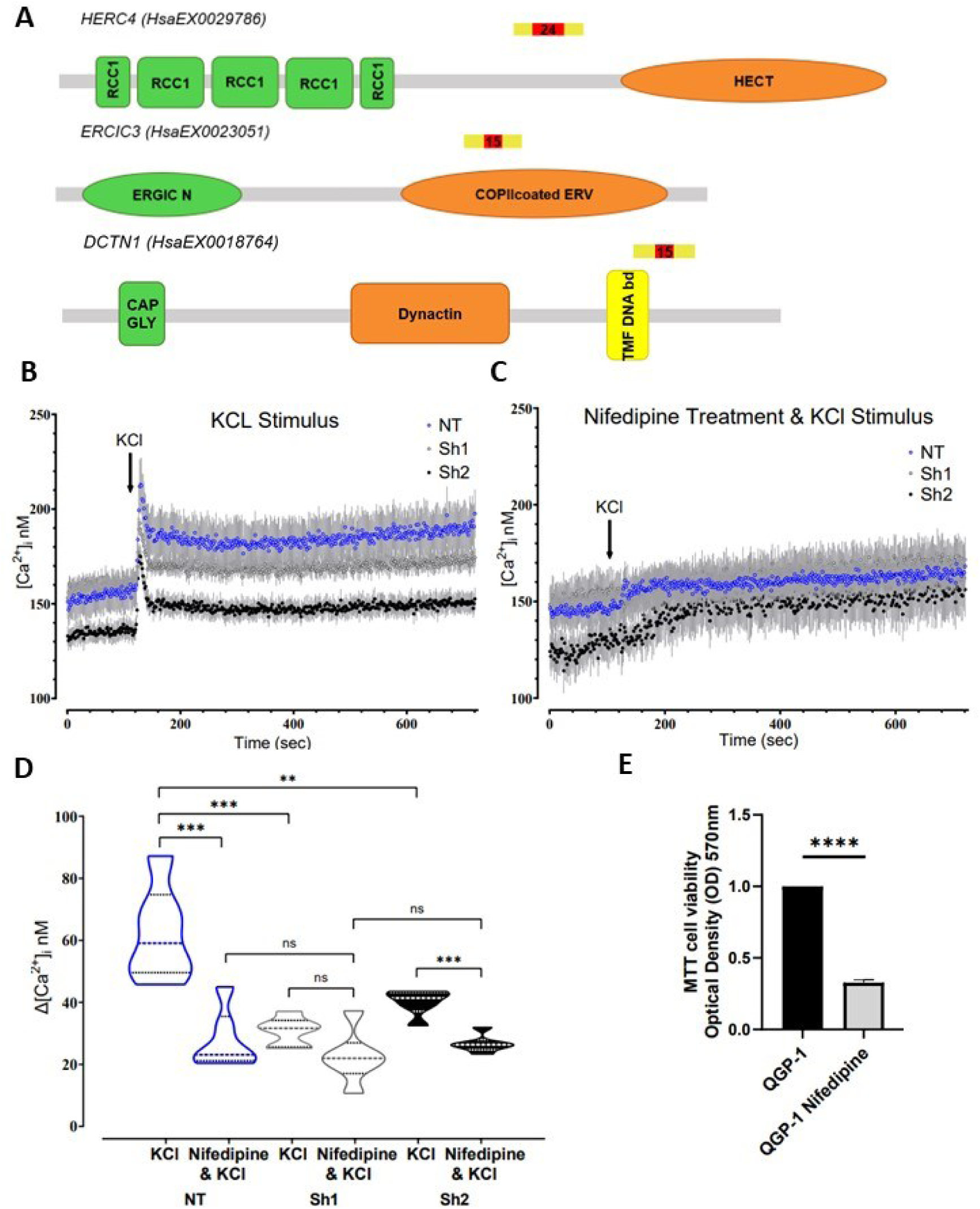
SRRM3 regulates intracellular calcium responses in PanNET cell lines. (**A**) Schematic showing the anticipated effect of alternative splicing events on the resulting proteins HERC4, ERGIC3, DCTN1. The position of the alternative exons in relation to the known functional domains are marked in each case. RCC1: Regulator of Chromosome Condensation 1, HECT: Homologous to the E6-AP Carboxyl Terminus domain, ERGIC N: Ergic N-terminal region, COPIIcoated ERV: COPII-coated endoplasmic reticulum-derived vesicles, CAP GLY: CAP-Gly domain, Dynactin, TMF DNA bd: TMF DNA-binding domain. (**B**) (**C**) [Ca^2+^]_i_ levels of QGP-1 NT or Sh1 or Sh2 cells. Error bars represent ±SEM. The arrows indicate the timepoint (120 sec) at which the stimulus (50 mM KCl) was added. In (**C**) nifedipine (10μM) was added in the wells 5 min prior to addition of KCl. (**D**) Violin plots summarize the differences in [Ca^2+^]_I_ (Δ [Ca^2+^]_i_) at the start and end of intracellular calcium measurements shown in B and C. Two-tailed unpaired t-tests were used to detect statistically significant changes.**, p<0.01; ***, p<0.001. (**D**) Bar-plot from MTT assay measured at 570nm illustrating the decreased viability in QGP-1 cells upon Nifedipine treatment. For statistically significant changes, two-tailed unpaired t-test was used.****, p<0.0001.

### *SRRM3* controls insulin granule exocytosis in PanNET cells

The reduced increase of intracellular Ca^2+^ levels upon SRRM3 downregulation in PanNET cells and the fact that Ca^2+^ influx through voltage-gated Ca^2+^ channels is essential for insulin exocytosis^45^ prompted us to assess the effect of *SRRM3* downregulation on secretory granule exocytosis in these cells. For this, we used the ratiometric reporter RINS1^26^, which is based on a double-fluorescent-fusion (superfolder GFP, sfGFP and mCherry) construct of proinsulin and allows imaging and quantification of insulin secretion. When cells transfected with this construct are stimulated by elevated glucose levels that generate increases in intracellular calcium levels, mature secretory granules fuse with the plasma membrane and release Insulin tagged with sfGFP (Ins-sfGFP), whereas the inactive C-peptide is tagged by mCherry and remains intracellular. We transfected with RINS1 either NT-3 cells treated with SRRM3 specific siRNAs or the control cells, treated with non-specific scrambled siRNAs and observed by confocal microscopy secretory granules before and after stimulation of the cells with 20mM glucose/30mM KCl (Figure 6A). More specifically, in unstimulated cells, we could detect the intracellular RINS1 reporter (colocalization of the two fluorescent signals) as expected. Upon stimulation of control cells, a clear increase of the intracellular mCherry signal and separation from the sfGFP one in the parental cells were observed after 10 minutes, indicating the presence of insulin-sfGFP in granules preparing to be released/and or released from the cells, while mCherry-Cpeptide remained in the cells. However, in *SRRM3* siRNA treated cells there was no clear separation between the two signals (mCherry and sfGFP) 10 minutes after stimulation with glucose/KCl and the cells were similar to the unstimulated ones, indicating problematic insulin granule maturation and exocytosis in the absence of *SRRM3.* Quantification of the fluorescent signals in the supernatants verified the overall reduced ability of the *SRRM3* KD cells to secrete insulin upon stimulation (Figure 6B). Furthermore, the difference in the secretion levels of insulin-sfGFP was confirmed by detecting insulin (sfGFP-Ins) in the cell supernatant by western blot analysis (Figure S6A), where reduced levels were identified to be secreted in cells with reduced levels of *SRRM3*.

**Figure 6.**
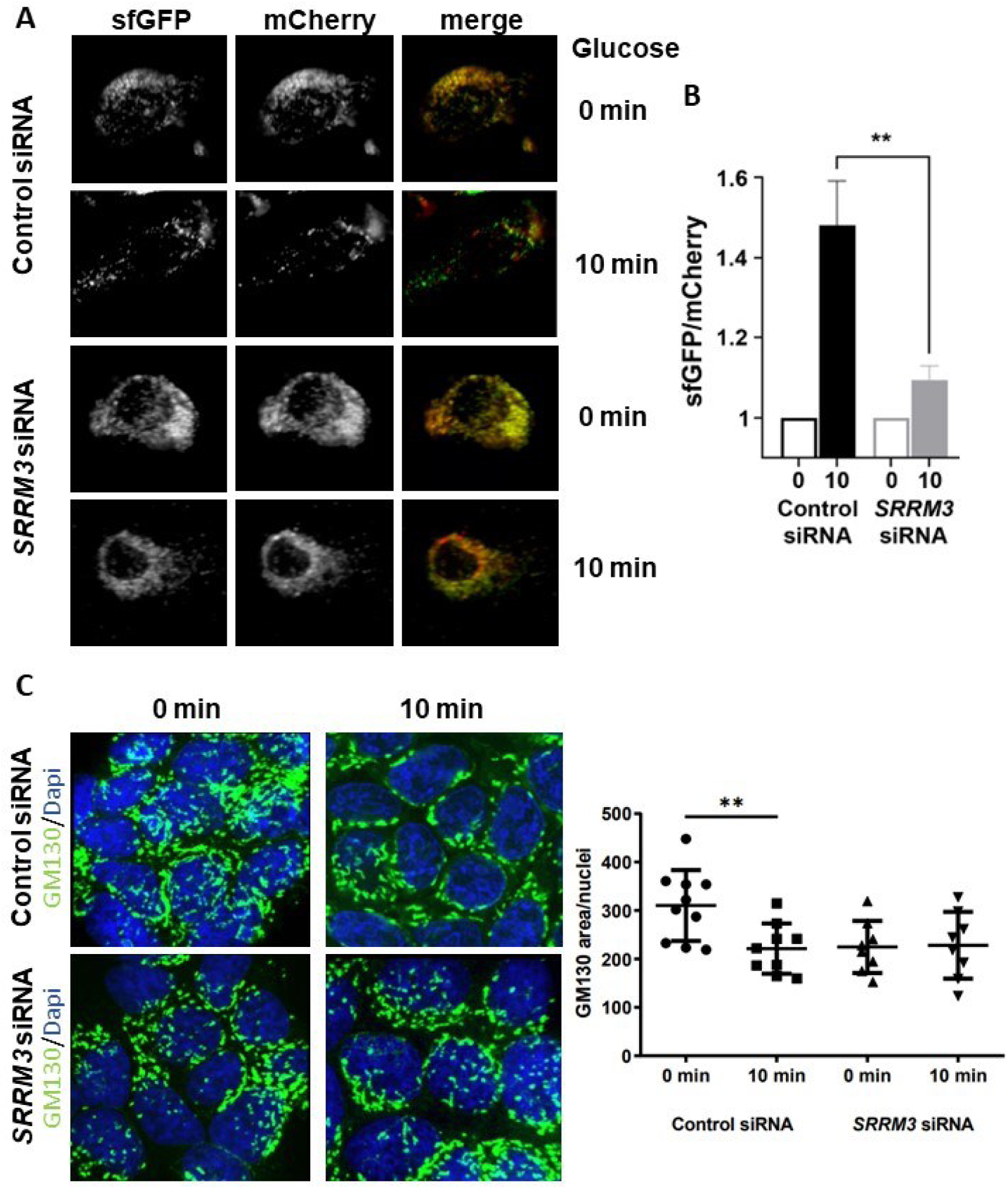
SRRM3 down-regulation impairs insulin secretion and alters Golgi morphology in PanNET cells. (A)-(B) NT-3 cells co-transfected with control or SRRM3 siRNA and pRINS1 were stimulated with 20 mM glucose/30 mM KCl for 0 and 10 min. (A) Representative confocal single plane images of cells presented in (B) to monitor the colocalization of sfGFP and mCherry in intracellular vesicles. (B) Insulin secretion was estimated by the ratio of the sfGFP versus the mCherry signal intensity in the supernatant, after normalization against the respective signals in the lysate. The graph shows the mean fold-change in the sfGFP/mCherry signal 10 min after stimulation relative to t0 of 3 independent experiments. Error bars represent ±SEM. **, p<0.01. (C)-(D) QGP-1 cells transfected with either control or SRRM3 siRNA, unstimulated or stimulated with 20mM glucose/30mM KCl for 10 min were stained with antibodies against GM130 Golgi marker and DAPI. (C) Representative confocal images of cells presented in (D) to monitor the cellular area occupied by Golgi relatively to nuclei area. (D) The bar graph presents the mean Golgi/nuclei area where ten cells from each category were used for the quantification. For statistically significant changes, two-tailed unpaired t-test was used.**, p<0.01.

**Figure S6.**
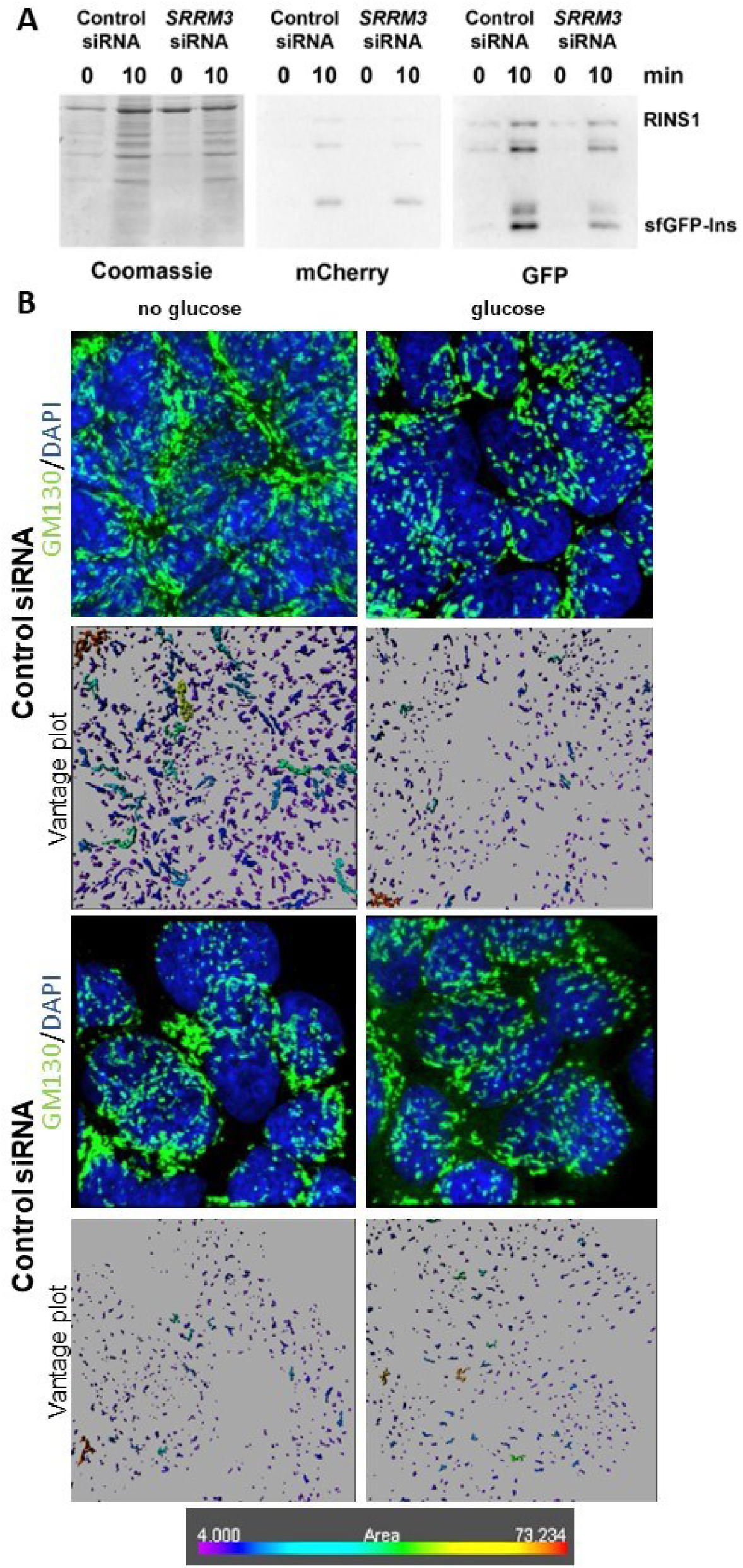
SRRM3 down-regulation impairs insulin secretion and alters Golgi morphology in PanNET cells. (**A**) QGP1 cells co-transfected with control or SRRM3 siRNA and pRINS1 were kept unstimulated or stimulated with 20 mM glucose/30 mM KCl for 10 min. Western blot analysis was performed to confirm insulin secretion in the supernatant samples presented in (Figure 6A, B). Anti-GFP blotting was used to detect the secreted processed insulin (sfGFP-Ins), which is not detected by anti-mCherry antibodies. Coomassie staining was used to monitor loading. (**B**) Confocal images of QGP-1 cells, transfected with control (scramble) or SRRM3-specific siRNAs, unstimulated (no glucose) or stimulated with 20 mM glucose/30 mM KCl (glucose) for 10 min and subsequently immunostained against GM130 (cis-Golgi) and DAPI. The Vantage plots underneath each image were generated using Imaris Microscopy Image Analysis software and show the range of the area covered by the cis-Golgi objects identified. The scale bar at the lower panel indicates the area size detected.

Knowing that the dynamic regulation of Golgi morphology is critical for the transport and packaging of secreted molecules into nascent secretory vesicles and for the progression and metastasis of cancers^46^ and given that many of the SRRM3 regulated microexons alter the function of components of the Golgi vesicle transport process (Figure 3D) we examined the morphology of Golgi upon SRRM3 downregulation and in response to stimulation by glucose. *SRRM3* downregulation resulted in reduction of the cellular area covered by Golgi (Figure 6C, D, S6B), changing the fragmentation level of Golgi. Furthermore, stimulation of the cells by glucose induced significant changes to the morphology of Golgi only in the presence of *SRRM3.* This finding supports the participation of SRRM3 in the process of glucose-induced vesicle mediated transport.

### SRRM3 supports PanNET growth in vitro and in vivo via regulation of a small number of microexons

To assay the importance of SRRM3 on tumour growth in PanNET, we firstly assayed the QGP-1-derived cell lines for proliferation. Both QGP-1-sh1 and sh2 showed reduced proliferation rates in the absence of SRRM3 compared to the control cells, QGP-1-NT (Figure 7A). Moreover, we assayed the ability of these cell lines to form colonies and detected the reduced ability for colony formation upon SRRM3 downregulation (Figure 7B). A change in the ability of QGP-1 cells to form spheres was also noted (Supplementary Figure S7A), in agreement with the dependency of these cells on SRRM3 for the development of PanNET phenotypic characteristics.

**Figure 7.**
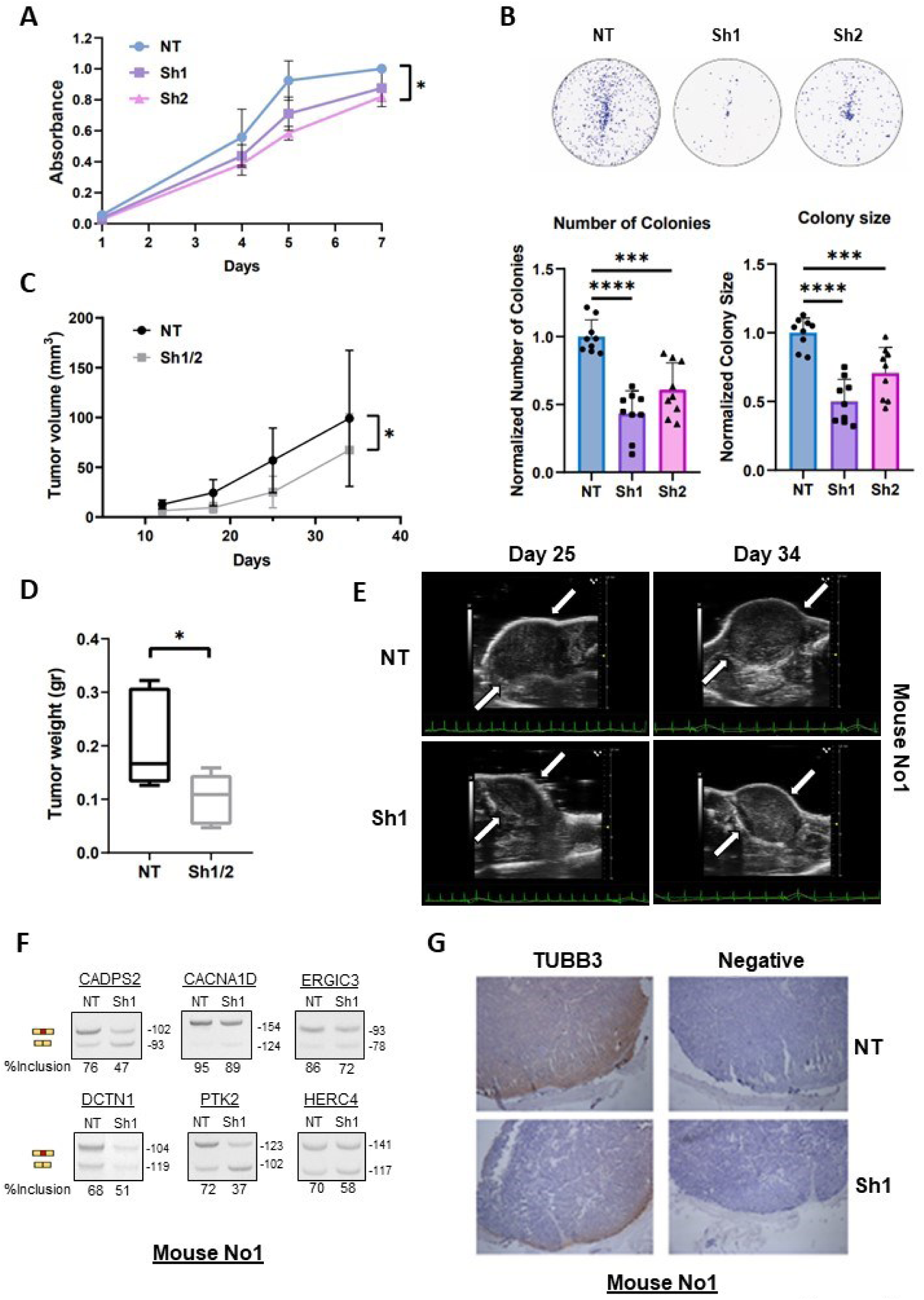
SRRM3 promotes tumor growth *in vitro* and *in vivo*. (**A**) Cell growth curve for QGP1 cells treated either with non-specific (NT), or with *SRRM3*-specific (Sh1, Sh2) shRNAs. (**B**) Colony formation assay of QGP1 NT, Sh1, Sh2 cells. Bar-plots of quantification of colony numbers and sizes are depicted as shown. Two-tailed unpaired t-test was used to detect statistically significant changes. ***, p<0.001, ****, p<0.0001. (**C**)-(**F**) QGP1 NT, Sh1, Sh2 cells were subcutaneously injected into the left (NT) and the right flank (Sh1/2) of NOD-SCID mice and tumors were monitored weekly over a period of 34 days. In (**C**), tumor volume was measured using VevoLAB software (v5.8.1.3266) and visualized using GraphPad Prism (v8), where results suggest a correlation between Sh cells and reduced tumor growth. Two-tailed paired t-test was used to detect statistically significant changes. *, p<0.05. In (**D**), graph of the tumor weights at end-point are shown *, p<0.05. In (**E**), 3D ultrasound images of mouse No1 are presented as an example of the last two timepoints of the experiment. Arrows are used to delineate the upper right and lower left corners of the tumors. Electrocardiograms (ECGs) are presented at the bottom of the images in green. Size scale bar is next to the figures in mm. (**F**) Analysis by RT-PCR and gel electrophoresis of the indicated splicing events in mouse No1. %inclusion values represent the quantification of the inclusion band in ratio with the total signal from the inclusion and skipping bands. Molecular lengths (bp) are marked on the right of each picture. (**G**) Sections of tumors from mouse No1, derived from both NT and Sh1 cells, were embedded in paraffin, sliced, and subjected to Hematoxylin staining. Additionally, they were probed with antibodies against TUBB3. Representative images of TUBB3 staining and negative controls are provided.

**Figure S7.**
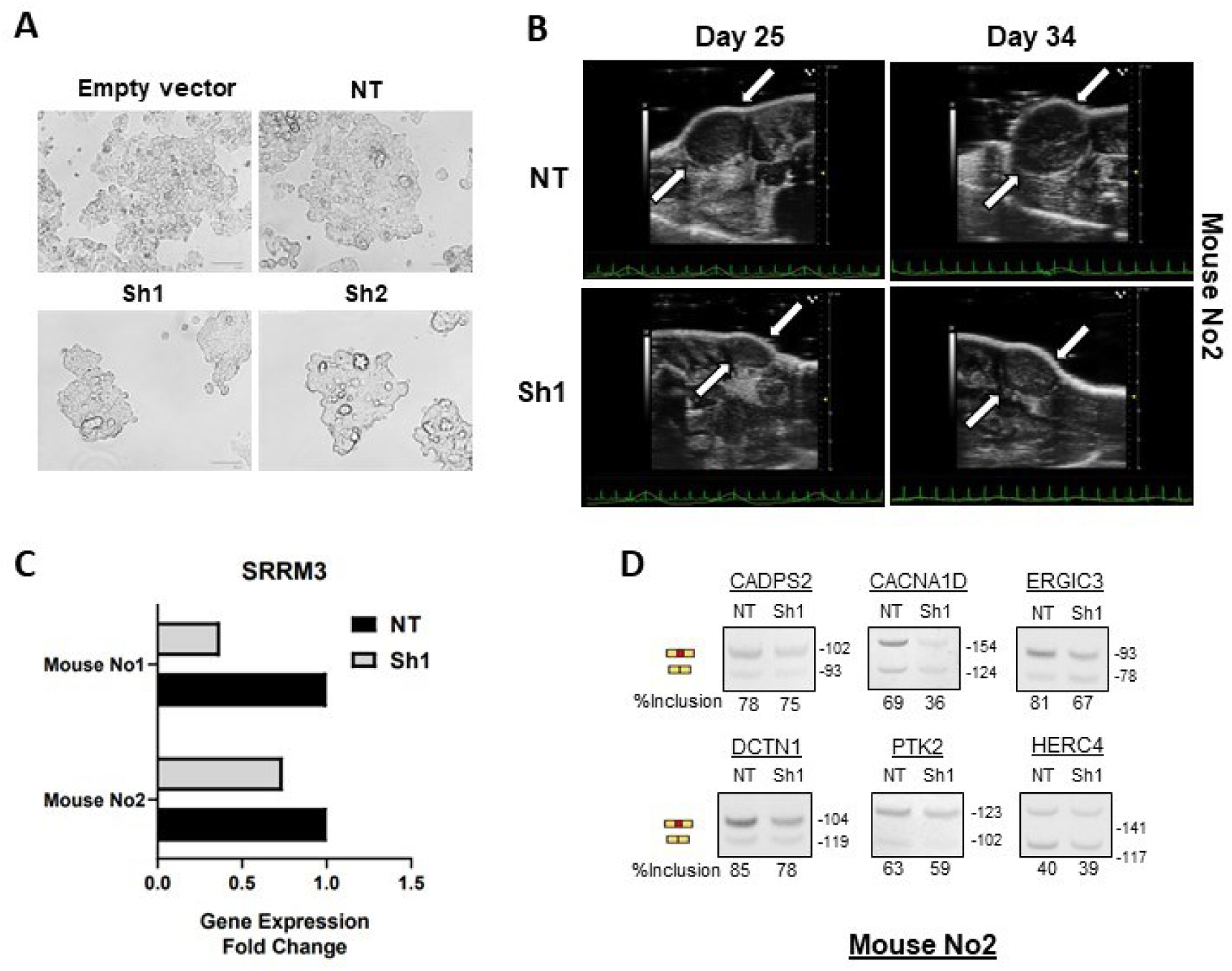
SRRM3 promotes tumor growth *in vitro* and *in vivo*. (**A**) Microscopy (phase-contrast) images of QGP1-cells treated with non-specific (NT) or *SRRM3*-specific shRNAs (Sh1, Sh2). (**B**) 3D ultrasound images of mouse No2 are presented as an example of the last two timepoints of the experiment. Arrows are used to delineate the upper right and lower left corners of the tumors. Electrocardiograms (ECGs) are presented at the bottom of the images in green. Size scale bar is next to the figures in mm. (**C**) Bar-plot of RT-qPCR analysis of *SRRM3* expression in tumors extracted from two mice from the xenograft experiment, to verify the downregulation of SRRM3 in the tumors derived from Sh cells in contrast to NT tumors. *GAPDH* levels were used for the normalization. (**D**) Analysis by RT-PCR and gel electrophoresis of the indicated splicing events in mouse No2. %inclusion values represent the quantification of the inclusion band in ratio with the total signal from the inclusion and skipping bands. Molecular lengths (bp) are marked on the right of each picture.

Subsequently, the SRRM3-KD and the respective control QGP-1-derived cell lines were used in xenotransplantation experiments in NOD-SCID mice. Tumour growth curves showed significant reduction in the progression of tumours derived from cells lacking *SRRM3* (Figure 7C). At the end point, significantly reduced tumour weight was measured as a result of *SRRM3* downregulation (Figure 7D). For accuracy and to keep the tumour volumes small we monitored tumour growth by ultrasonography during the course of the experiment (Figures 7E, S7B show representative images). At the end point, RNAs from tumours were isolated and used to verify the reduced levels of *SRRM3* and the expected effect on the inclusion of the microexons regulated by SRRM3 (examples in Figures 7F, S7C, S7D).

Interestingly, the fact that all the microexons that are deregulated in PanNETs depict increased inclusion levels specifically in neuronal tissues^15,19^, together with the overexpression of neuronal splicing factors suggests the prevalence of a neuronal phenotype in PanNETs. Given that tumour innervation affects the acquisition of various hallmark capabilities, such as increased proliferation and invasion^47^, we assessed the involvement of SRRM3 in tumour innervation. For this, we assayed the levels of the neuronal marker TUBB3 in the xenograft tumours derived from QGP1 or the generated SRRM3 knock-down cells (Figure 7G). We detected significant reduction in the staining intensity for the tumours that were derived from cells with reduced levels of SRRM3 (Figure 7G), strengthening our finding that SRRM3 is important for the development of a neuronal phenotype in PanNETs.

To verify that the effect of SRRM3 is via its ability to regulate the inclusion of neural microexons, we tested whether deregulation of individual microexons was sufficient to alter the PanNET characteristics that depend on SRRM3. For this, we designed antisense oligonucleotides (ASOs) harbouring 2’-O-methyl phosphorothioate modifications targeting microexon 3’ splice sites and adjacent regulatory sequences, including the UGC motif required for SRRM3-dependent control of microexon inclusion^32^. We initially selected the microexons of *CADPS2*, *PTK2* and *CACNA1D* as the most deregulated microexon events among the validated ones and because they affect critical components of calcium-induced secretory pathways (Figure 8). In all three cases introduction of the ASOs in QGP1 cells, in comparison to control siRNAs, induced microexon skipping, as expected. The ASOS against the *CACNA1D* and *PTK2* microexons were more efficient in inducing microexon skipping to levels similar to the *SRRM3* downregulation (Figure 8A), however, using a pool of the three ASOs resulted in a better skipping for all three microexons (Figure 8B). Following this, we tested the effect on PanNET characteristics. It was prominent that upon treatment with the pool of ASOs, the formation of the characteristic PanNET cell spheres was reduced (Figure 8C). At the same time, the levels of the neuronal marker TUBB3 were significantly lower, as it was detected by immunofluorescence confocal microscopy (Figure 8D). Using MTT assay, 72 hours after transfection, we tested whether the increased skipping induced by the ASOs affected the viability of the QGP1-cells. As shown in Figure 8E a significant reduction of the viability of the PanNET cells was detected as a result of the increased skipping of the microexons of *CADPS2, CACNA1D* and *PTK2.* Taken together, these results point to SRRM3 and the neural microexons regulated by SRRM3 as molecules that could be investigated for the development of PanNET therapeutic approaches.

**Figure 8.**
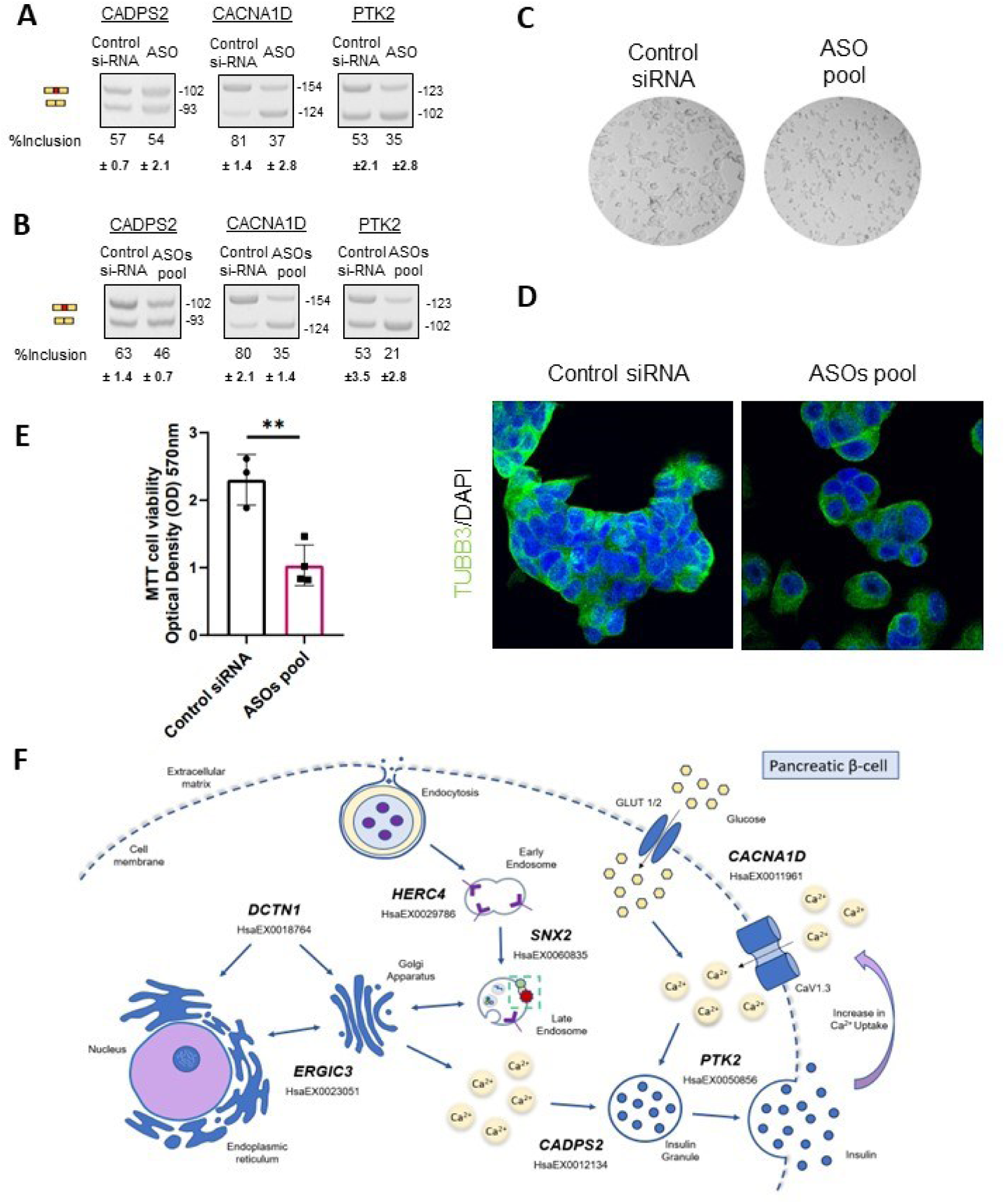
Increased skipping of a set of SRRM3-regulated microexons alters PanNET cell characteristics. (**A**) Analysis by RT-PCR and gel electrophoresis of the indicated splicing events in RNA samples derived from QGP-1 control cells (treated with scramble siRNAs) and QGP-1 treated with specific ASO for each splicing event. %inclusion values represent the quantification of the inclusion band in ratio with the total signal from the inclusion and skipping bands. Molecular lengths (bp) are marked on the right of each picture. (**B**) RT-PCR analysis of the indicated events in RNA samples derived from QGP-1 control cells (treated with scramble siRNAs) and QGP-1 treated with a pool of the three ASOs presented in A. The %inclusion values represent the quantification of the inclusion band relative to the total signal from both the inclusion and skipping bands. Error bars, based on the standard deviation (±SD) values from the corresponding biological replicates, are depicted below. Product lengths (bp) are marked on the right of each picture. (**C**) Microscopy (phase-contrast) images of QGP1-cells treated with control (scramble) siRNA or pool of the three ASOs. (**D**) Confocal images of QGP-1 cells transfected with control siRNA or pool of the three ASOs after immunostaining against TUBB3. (**E**) Bar-plot from MTT assay measured at 570nm illustrating the decreased viability in QGP-1 cells 72h after transfection with pool of ASOs in comparison to control (scramble) siRNAs. For statistically significant changes, two-tailed unpaired t-test was used.**, p<0.01. (**F**) Schematic representation of the secretory pathways that promote PanNET growth through cancer-favoring inclusion of neural microexons in the indicated mRNAs, all regulated by the splicing factor SRRM3. The VastDB ID numbers of the events are shown underneath the names of the mRNAs.

## Discussion

In the present study, we analysed data from a large number of PanNET patient samples aiming to identify AS-derived isoforms that characterize PanNETs. We show that the deregulation of AS in PanNETs results in the development of a neuronal phenotype in these tumours, with the overexpression of a number of neuron-specific splicing factors, and the subsequent upregulation of a neural splicing program. We pinpoint to the regulatory splicing factor *SRRM3* as an unexplored source of this phenotype that controls the inclusion levels of a number of microexons in PanNETs, the majority of which have been characterized as neural microexons^19^. These microexons code for surface-accessible, charged amino acid stretches that overlap or lie near functional protein domains, thus contributing to the remodelling of protein-interaction networks that operate during neuronal maturation^19^.

Interestingly, in PanNETs, deregulation of microexons, driven by the overexpression of SRRM3, results in alteration of the secretory abilities of neuroendocrine cells. Most of the altered microexons in PanNETs display high inclusion levels in pre-mRNAs associated with axon development, regulation of trans-synaptic signalling and vesicle mediated transport and exocytosis. Involved in such functions, a 9nt neuronal alternative microexon incorporated in the mRNA of CADPS2 alters a domain responsible for the interaction of this protein with the dopamine receptor DRD2. *CADPS2* is a gene coding for a secretory granule associate protein that mediates monoamine transmission and neurotrophin release. Alternatively spliced variants of the *CADPS2* mRNA have been connected to autism spectrum disorder^48^ and are subjected to further analyses on their involvement in neurodegenerative diseases^49^. The increased inclusion of a neuronal microexon in *CACNA1D* pre-mRNA affects the ability of the L-type voltage gated Ca^2+^ channel Cav1.3 to regulate intracellular calcium levels (this study), affecting not only the trafficking of secretory vesicles^50^, but also the viability of PanNET cells (this study). Similarly, the upregulated microexon of PTK2 results in a protein isoform that is the main isoform in forebrain neurons^42^, depicts increased autophosphorylation levels, decreased phosphorylation by Src thereby potentially affecting insulin granule exocytosis^43,51^, Golgi structure and integrity^52^. Similarly, increased inclusion of the neuronal alternative microexon of *ERGIC3* pre-mRNA inserts 5 amino acids at the COPII-ERV domain, potentially affecting the functionality of ERGIC3 in processing in the ER and/or sorting of molecules into COPII vesicles for transport to the Golgi^53^. *HERC4* (HECT Domain and RCC1-Like Domain 4) encodes a protein classified as a HECT and RLD (RCC1-like domain) containing E3 ubiquitin protein ligase. HERC4 seems to be involved in protein trafficking, the distribution of cellular structures and cellular organization^54^. HERC4 interacts with USP8 protein, which regulates the morphology of the endosome and is involved in cargo sorting and membrane trafficking at the early endosome stage^55^. *DCTN1* codes for Dynactin subunit 1, the largest subunit of the dynactin complex, an activator of the molecular motor protein complex dynein^56^. Plays a key role in dynein-mediated retrograde transport of vesicles and organelles along microtubules^57^. Independently of dynein-mediated axonal transport, DCTN1 protein promotes motor neuron synapse stability^56^.

The majority of the microexons deregulated in PanNETs attest increased inclusion in both functional and non-functional types of the disease. At least partially, this deregulation is driven by the altered expression of SRRM3, which is similar in functional and non-functional tumours and is increased with disease progression. Thus, it is not straightforward to assume that overexpression of SRRM3 alters the ability of neuroendocrine cells to produce hormones, as its role, at least in PanNETs, is probably more complicated. The fact that SRRM3 downregulation significantly affects the ability of PanNET cells to proliferate, *in vivo* and *in vitro* suggests that its role extends beyond hormone secretion. It is known that cancer cells release diverse neurotrophic factors and exosomes to promote tumour innervation^47,58^. A growing body of evidence suggests that cancer prognosis is related to intratumoural nerve infiltration, especially in organs like pancreas, which displays high innervation^47,59^. The density of infiltrated nerves is positively associated with tumour metastasis, morbidity, and mortality. Furthermore, innervation of cancers apparently plays an important direct role in promoting metastasis. On the other hand, vesicle mediated trafficking and endocytosis and exocytosis are important mechanisms facilitating intratumoural cell communication^46,60^. Our results suggest that SRRM3 and the regulated-microexons have a significant role in the majority of the above-described cellular processes and could be regarded as targets for the development or repurposing of drugs targeting tumour innervation and secretion.

## Conclusion

Concluding, we have shown that a neural splicing factor, SRRM3, is a master regulator of Alternative Splicing in Pancreatic Neuroendocrine Tumors (PanNETs). SRRM3 is upregulated in PanNETs and controls the inclusion of a small group of microexons, that in normal tissues are included only in mRNA isoforms specific for neural cells. As a result, the secretory function of pancreatic islet cells is altered in many levels (response of intracellular calcium to stimulus, vesicle mediated transport, Golgi morphology) and this promotes cancer cell traits. We have also shown that by targeting a group of three SRRM3-controlled microexons can dramatically change the ability of cancer cells to proliferate.

## Declarations Ethical Approval

Animal studies were approved by the Institutional Committee of Protocol Evaluation in conjunction with the Veterinary Service Management of the Hellenic Republic Prefecture of Attika according to all current European and national legislation under the license: 278206-01/04/2022. Experiments were performed in accordance with the guidance of the Institutional Animal Care and Use Committee of BSRC “Alexander Fleming’.

## Authors’ contributions

M.P. performed the bioinformatics analyses and performed or supervised cellular assays, animal studies and the in vitro splicing analyses, generated figures and contributed in writing the manuscript. C.M. performed the majority of the splicing analyses, contributed in the animal and cellular studies. Z.E performed the RINS1 experiments and prepared Figures 6A, B and S6A. G.R. and V.K. advised on image acquisition and analyses and prepared Figures 6C and S6B. A.K. performed performed RNA preparations from animals and cells and islet isolation from mouse pancreas. A.M. performed the xenograft animal studies. M. D and V.N. performed the ultrasound imaging of the xenograft tumors and prepared Figures 7E, S7B. J.S. provided the PanNET cell lines and adviced on performing the cellular assays. J.J-M. provided advice on islet experimentation, provided ASOs for splicing experiments. S.G.D. advised and contributed in the calcium experiments and in preparation of Figures 5, S5. M.E.R performed part of the bioinformatics analyses and cellular assays, organized the bioinformatics analyses, supervised M.P. on the bioinformatics analyses. PK conceived, designed and supervised the study, wrote the manuscript text and prepared and supervised preparation of the Figures. All authors reviewed the manuscript.

## Availability of data and materials

All datasets used in the present study are publicly available as follows: PanNET public RNA sequencing datasets can be found under accession numbers EGAS00001001732 (Scarpa A., et al. Whole-Genome Landscape of Pancreatic Neuroendocrine Tumours. Nature. 2017;543:65–71. doi: 10.1038/nature21063) and GSE118014 (Chan, C. S. et al. ATRX, DAXX or MEN1 mutant pancreatic neuroendocrine tumors are a distinct alpha-cell signature subgroup. Nat Commun 9, (2018), doi: 10.1038/s41467-018-06498-2). Normal pancreatic tissue datasets derived from VastDB repository for each different type of pancreatic cell can be found at the following URL: https://vastdb.crg.eu/wiki/Downloads#Homo_sapiens_.28hg38.29. Also, the dataset for whole pancreatic islets can be found under accession number GSE50398 (Tapial, J. et al. An atlas of alternative splicing profiles and functional associations reveals new regulatory programs and genes that simultaneously express multiple major isoforms. Genome Res. 27, 1759–1768 (2017). doi: 10.1101/gr.220962.117). The human adenocarcinoma dataset can be found under accession number GSE79668 (Kirby, M. K. et al. RNA sequencing of pancreatic adenocarcinoma tumors yields novel expression patterns associated with long-term survival and reveals a role for ANGPTL4. Molecular Oncology 10, 1169–1182 (2016), doi: 10.1016/j.molonc.2016.05.004).

## Acknowledgments

We thank A. Dimas, V. Koliaraki, M. Tsoumakidou, G.K. Kollias (B.S.R.C. “Al. Fleming”), D.L. Kontoyiannis, (Aristotle University of Thessaloniki, Greece), J. Valcarcel, M. Irimia (C.R.G. Barcelona, Spain) for reagents. We would also like to thank Sophie Bonnal for help with the design of the ASOs.

