## Supplemental Table 5 for "A neural alternative splicing program controls cellular function and growth in Pancreatic Neuroendocrine tumours"

**RT-PCR Primers and Conditions**

HUMAN

|  | **Gene** | **Event ID** | **Primer Sequence (5’-3’)** | **Annealing Temperature (Tm)** | **GC Content** | **Amplicon length(bp)** | **Product**  **Size (bp)** |
| --- | --- | --- | --- | --- | --- | --- | --- |
| 1. | **CACNA1D** | EX0011961 | **F:**TTCTTCCCACAAGCATGACCG | 63 | 52 | 21 | 124-154 |
|  |  |  | **R:**GGGGTCCCTGAAATAGCCATG |  | 57 | 21 |  |
| 2. | **CADPS2** | EX0012134 | **F:**TCTTCATCTGGCAGAGCTCTG | 61 | 52 | 21 | 93-102 |
|  |  |  | **R:**TCAGCCAATAAATCAGGCCACC |  | 50 | 22 |  |
| 3. | **DCTN1** | EX0018764 | **F:**TTGCTACTCTGGTCTCTGGCA | 58 | 52.4 | 21 | 104-119 |
|  |  |  | **R:**CTGAAGCAGCAGTGGTGAGTC |  | 52.4 | 21 |  |
| 4. | **ERGIC3** | EX0023051 | **F:**AAACTTCCACTTTGCCCCTGG | 60 | 52.4 | 21 | 78-93 |
|  |  |  | **R:**TCAAGGCCAAAGCTCTGCAAG |  | 52.4 | 21 |  |
| 5. | **HERC4** | EX0029786 | **F:**CTGGGTCCAGCAGCAAGC | 60 | 67 | 18 | 117-141 |
|  |  |  | **R:**TGCATCTGTCTGGAGCAGAGT |  | 52 | 21 |  |
| 6. | **PTK2** | EX0050856 | **F:**TGAAGAAGATACTTACACCATGCCC | 61 | 44 | 25 | 102-123 |
|  |  |  | **R:**ACATCTCCAAATTGGCCTTCTCC |  | 48 | 23 |  |

MOUSE

|  | **Gene** | **Event ID** | **Primer Sequence (5’-3’)** | **Annealing Temperature (Tm)** | **GC Content** | **Amplicon length(bp)** | **Product**  **Size (bp)** |
| --- | --- | --- | --- | --- | --- | --- | --- |
| 1. | **CACNA1D** | EX0008762 | **F:**AGCATGTGTCTGAAAATGGGCA | 61 | 45 | 22 | 118-145 |
|  |  |  | **R:**TCTTCACGGCAAATGGTTGGA |  | 48 | 21 |  |
| 2. | **CADPS2** | EX0008897 | **F:**GCTCTGCATAGAAGTCCTACAGC | 58 | 52 | 23 | 99-108 |
|  |  |  | **R:**GCCCAAAACTTCTCTGCATGC |  | 52 | 21 |  |
| 3. | **DCTN1** | Ex0013951 | **F:**AGAGCTCAAGCAGCGCCTGAAC | 57 | 59.1 | 22 | 172-187 |
|  |  |  | **R:**CAGCAGTGGGGAGTCCTTCA |  | 60 | 20 |  |
| 4. | **ERGIC3** | EX0017296 | **F:**AAACTTCCACTTTGCCCCTGG | 61 | 52.4 | 21 | 78-93 |
|  |  |  | **R:**TCAAGGCCAAAGCTCTGCAAG |  | 52.4 | 21 |  |
| 5. | **HERC4** | EX0022805 | **F:**CTGGGTCCAGCAGCAAGC | 61 | 67 | 18 | 112-136 |
|  |  |  | **R:**TGCATCTGTCTGGAGCAGAGT |  | 52 | 21 |  |
| 6. | **PTK2** | EX0037737 | **F:**GACACATACACCATGCCCTCG | 62 | 57 | 21 | 117-138 |
|  |  |  | **R:**GCTCAGGTACACGCCTTGATG |  | 57 | 21 |  |

**qPCR Primers and Conditions**

MOUSE

|  | **Gene** | **Primer Sequence (5’-3’)** | **Annealing Temperature (Tm)** | **GC Content** | **Amplicon length(bp)** |
| --- | --- | --- | --- | --- | --- |
| 1. | **GAPDH** | **F:**TGCACCACCAACTGCTTAG | 59-60 | 52.6 | 19 |
|  |  | **R:**GGATGCAGGGATGATGTTC |  | 52.6 | 19 |
| 2. | **GCG** | **F:**CTTCCCAGAAGAAGTCGCCA | 59 | 55 | 20 |
|  |  | **R:**GATGAAGTCCCTGGTGGCAA |  | 55 | 20 |
| 3. | **HPRT** | **F:**GCTGGTGAAAAGGACCTCT | 59 | 53 | 19 |
|  |  | **R:**CACAGGACTAGAACACCTGC |  | 55 | 20 |
| 4. | **INS1** | **F:**TGTTGGTGCACTTCCTACCC | 59 | 55 | 20 |
|  |  | **R:**CTTCCTCCCAGCTCCAGTTG |  | 60 | 20 |
| 5. | **INS2** | **F:**CACCCAGGCTTTTGTCAAGC | 59 | 55 | 20 |
|  |  | **R:**CGGGACATGGGTGTGTAGAA |  | 55 | 20 |
| 6. | **SRRM3** | **F:**GAGGAGCCGCAAAAAGAGGAG | 60 | 57.1 | 21 |
|  |  | **R:**GACCGATCTCGACGGTGTTTC |  | 57.1 | 21 |

HUMAN

|  | **Gene** | **Primer Sequence (5’-3’)** | **Annealing Temperature (Tm)** | **GC Content** | **Amplicon length(bp)** |
| --- | --- | --- | --- | --- | --- |
| 1. | **GAPDH** | **F:**ACATCAAGAAGGTGGTGAAGCAGG | 60 | 50 | 24 |
|  |  | **R:**TGTCGCTGTTGAAGTCAGAGGAGA |  | 50 | 24 |
| 2. | **SRRM3** | **F:**CATCCAGTGGACAAGGAGAAG | 60 | 52 | 21 |
|  |  | **R:**GAGCTAAGGTGAAGTGACAGAG |  | 50 | 22 |
